# A non-canonical glucose metabolic rewiring enables resistance to cancer targeted therapy

**DOI:** 10.1101/2024.12.19.629335

**Authors:** Huimin Lei, Ruixue Xia, Yabin Tang, Jun Lu, Chan Xiang, Yujing Li, Hongyu Liu, Peichen Zou, Ayinazhaer Aihemaiti, Wei Zhang, Ying Shen, Baohui Han, Yuchen Han, Hua Zhong, Hongzhuan Chen, Liang Zhu

**Author notes:** Corresponding author. (L.Z.); (H.C.); (H.Z.). These authors contributed equally to this work (Huimin Lei, Ruixue Xia, Yabin Tang, Jun Lu).

## Abstract

Targeting glucose metabolism for cancer therapy is a long-lasting pursuit but with disappointing translational success. Revelation of actionable distinctions between the malignancy traits to achieve selectivity is a prerequisite for fulfilling a therapeutic window. We discover a non-canonical glucose transporter GLUT6-facilitated, lactate-independent, glucose metabolic rewiring that selectively enables resistance to targeted therapy in lung cancer. Downstream, GLUT6 facilitated glucose influx and straying to methylglyoxal which dimerized KEAP1 and upregulated NRF2 pathway to confer resistance. Upstream, GLUT6 was transcriptionally upregulated by therapy-induced MAZ activation. Targeting GLUT6 prevented and overcame resistance to EGFR and KRAS inhibitors and the MAZ-GLUT6-NRF2 signaling correlated with clinical therapy response and relapse. Our findings uncover an unrecognized non-genetic, metabolic mechanism for drug resistance and a scenario specifically determined by a distinct glucose metabolic rewiring in cancer. The preferential dependance of the non-canonical, general-homeostasis-less-perturbing transporter GLUT6 implies a promise for resistance overcoming and for glucose metabolism-targeting strategy, two long and urgent quests.

## Introduction

The resurgence of the wisdom that metabolic reprogramming characterizes and determines tumor malignancy highlights one of the recent cancer research advances ^1–3^. Cancer cells rewire metabolism to satisfy their aberrantly bioenergetic and biosynthetic demands ^2, 3^. Targeting the rewired metabolism is considered a promising anti-cancer strategy ^1–3^.

Glucose, the principal nutrient for cells, is the earliest metabolite found gone awry in cancer metabolism, inspiring enthusiasm aiming the deviant metabolism as the primary target for cancer therapy ^3–5^. However, this strategy has never been therapeutically successful in clinic practice though long-lasting pursuit, mainly due to the lack of understanding of the actionable distinctions between the malignancy traits to achieve enough therapeutic window ^1, 3, 5^. Therefore, the discovery and mechanism revelation of the distinctions will help to provide therapeutic margin capable of specifically hitting the cancer cells addicted to the metabolism, making glucose rewiring in this context a promising anti-cancer target.

Here we discover that GLUT6, a non-canonical Class III glucose transporter reported least affecting general metabolism, and the GLUT6-facilitated, lactate-independent, glucose metabolic rewiring enable targeted therapy resistance in lung cancer, promising a potential therapeutic margin for glucose metabolism-based cancer interception. This finding uncovers a novel diagram of glucose metabolism selectively demonstrated in and underlying cancer targeted therapy resistance, not only donating a chance to overcome the resistance, the principal challenge and clinical urgently unmet need ^6, 7^, but also implying a promise to target glucose metabolism more selectively, a long-lasting quest of strategy in cancer science ^1, 3^.

## Results

### Additionally exacerbated glucose uptake and addiction in therapy-resistant tumors

Cancer cells are characterized by increased glucose uptake and utilization which can be calibrated through ^18^F-fluorodeoxyglucose (FDG)-based positron emission tomography (PET) functional imaging analysis of the maximum standardized uptake value (SUVmax) ^3, 8^. PET-computer tomography (CT) analysis demonstrated a positive tumor SUVmax signal and an additional increase of SUVmax was noticed by us when the tumor relapsed to the epidermal growth factor receptor (EGFR) inhibitor (EGFRi) treatment of the EGFR mutant lung cancer (Figure 1A left panel, 1B), a paradigm of molecularly targeted therapy in precise medicine era ^9–11^.

**Figure 1.**
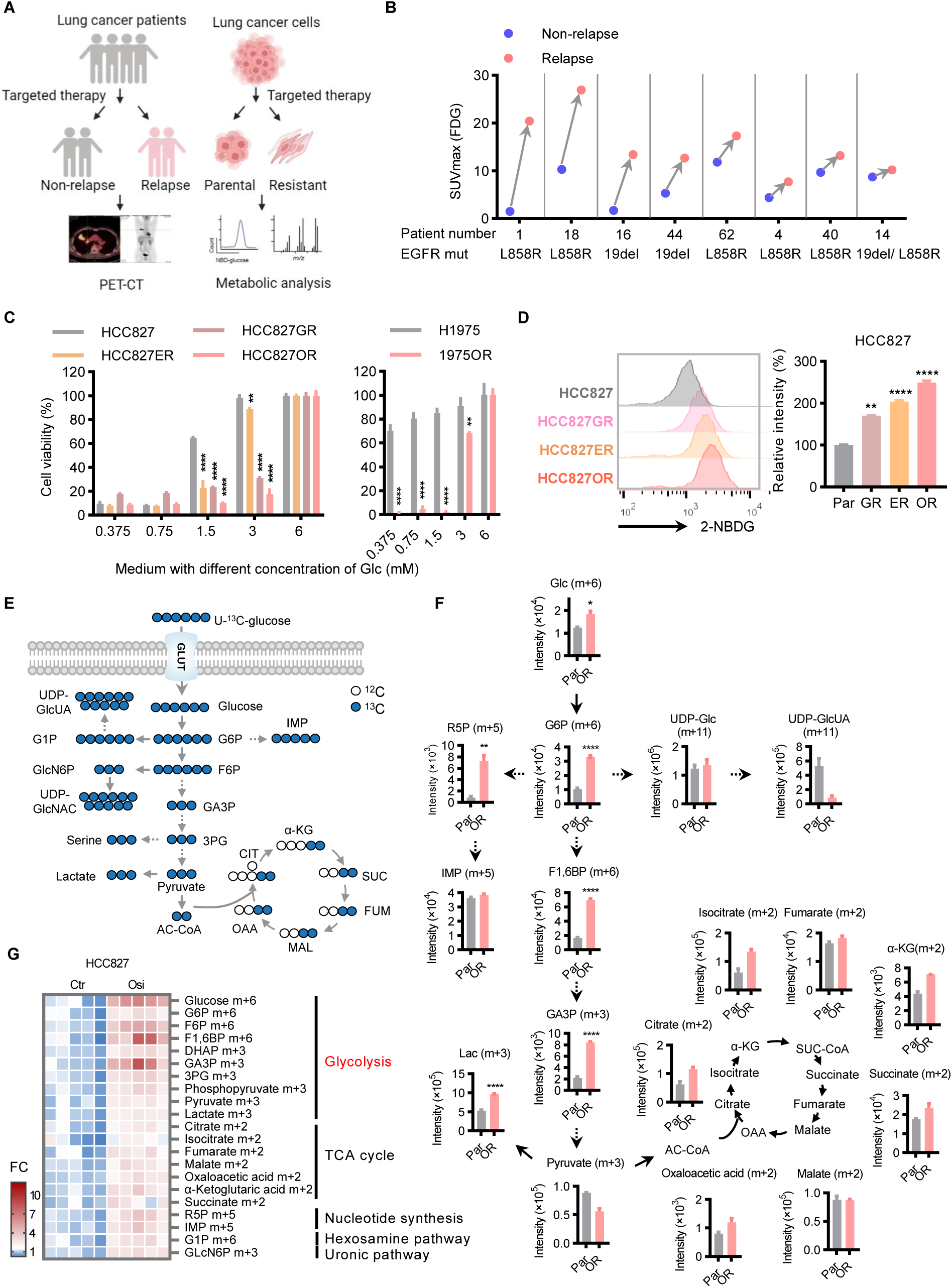
Additionally exacerbated glucose uptake and addiction in therapy-resistant tumors. (A) Schematic overview of glucose uptake analysis in clinical patients measured by ^18^F-FDG-PET-CT (left panel) and metabolic analysis in cell line models resistant to targeted therapy (right panel). FDG, fluorodeoxyglucose; PET, positron emission tomography; CT, computer tomography. (B) Increase of tumor maximum standardized uptake value (SUVmax) measured by PET-CT analysis at treatment-relapse phase versus paired the same person non-relapse (response and pre-treatment) phase in patients (n=8) underwent EGFRi targeted therapy. EGFR mutation status are shown. (C) More vulnerable to decreased abundance of medium glucose for viability in resistant versus parental cells. ER, erlotinib resistant; GR, gefitinib resistant; OR, osimertinib resistant. Cells were cultured in the medium with indicated concentrations of glucose (Glc) for 72 h. (D) Enhanced intracellular accumulation of fluorescent glucose analog 2-(N-(7-nitrobenz-2-oxa-1,3-diazol-4-yl) amino)-2-deoxyglucose (2-NBDG) in resistant versus parental cell assayed by using flow cytometry after the cells were cultured in glucose-free medium containing 100 μg/mL 2-NBDG for 30 min. (E) Schematic overview illustrating tracing of medium U-^13^C-glucose uptaken into cells and funneled into canonical metabolic pathways, the glycolysis, tricarboxylic acid (TCA) cycle, pentose phosphate, hexosamine, and serine biosynthesis. The metabolites and the number of labeled carbons were indicated in color. (F) Enhancement of labeled glucose uptaken and funneled into glycolysis in resistant cells versus parental cells. Cells were cultured in medium where glucose was replaced with the same concentration (11 mM) of U-^13^C-glucose for 6 h. The labeled metabolites were determined by LC/MS-MS-based stable isotope– resolved metabolic analysis (SIRM) analysis. (G) Intracellular accumulation of uptaken glucose and of glycolytic intermediates under osimertinib acute treatment (1 µM for 6 h) assayed by SIRM analysis. For (C), (D), and (F), (G) data are representative of at least three independent experiments with three to five technical replicates and presented as means ± SEM. *, *P* < 0.05; **, *P* < 0.01; ***, *P* < 0.001; ****, *P* < 0.0001.

In line with the additionally exacerbated glucose uptake and utilization in clinically targeted therapy resistant tumors, a pronouncedly preferred glucose requirement was identified in EGFRi-resistant cell lines compared with their isogenic sensitive parental counterparts (Figure 1A right panel, 1C, S1A-S1C). These clinically-relevant ^12, 13^ (Figure S1D) resistant cell line models were authenticated in this study (Figure S1E) and in our previously reported ^14–17^. When the medium glucose concentration decreased, the resistant cells exhibited much more vulnerable than parental cells in viability (Figure 1C, S1A), colony formation (Figure S1B), and growth (Figure S1C) analyses. In contrast to their addiction to glucose, the resistant cells compared with parental counterparts did not show more reliance on glutamine or serum in culture medium (Figure S1A), suggesting a selective dependence of the resistance on glucose supplementation.

The enhancement of glucose uptake in resistant cells was validated in analysis of transmembrane transportation of fluorescent glucose analog 2-NBDG [2-(N-(7-nitrobenz-2-oxa-1,3-diazol-4-yl) amino)-2-deoxyglucose]) (Figure 1D, S2A). This finding was orthogonally reconfirmed by liquid chromatography coupled to tandem mass spectrometry (LC-MS/MS) tracing analysis (Figure 1E) of the stable isotope ^13^C-glucose replacing the culture medium glucose at the same concentration, as shown by an intracellular abundance increase of the labeled glucose and its downstream glycolytic intermediates in resistant versus parental counterpart cells (Figure 1F, S2B, S2C).

The stable isotope–resolved metabolic analysis (SIRM) of ^13^C-glucose further demonstrated that the enhanced glucose metabolism in resistant cells compared with parental cells was apt to culminate in glycolysis, showing a preferred accumulation of glucose-derived, labeled intermediate metabolites in glycolytic pathway than in other major pathways (Figure 1E, 1F). The live-cell extracellular acidification rate (ECAR) analysis validated the enhanced glycolysis in resistant versus parental cells (Figure S2D, S2E). This glycolysis proclivity represents a reminiscence of cancer cell response to drug acute insult where the sensitive cells demonstrated an increase of intracellular uptaken glucose and the glucose-derived glycolytic intermediates when treated by osimertinib for 6 hours (Figure 1G).

These clinical patient and in vitro cell model data indicate that the targeted therapy-resistant cancers are characterized by an additional enhancement of glucose uptake, glucose addiction, and glycolysis compared with the sensitive ones.

### The rewired glucose uptake and metabolism in resistant cells are determined by a non-canonical transporter GLUT6

Glucose cellular uptake and utilization is rate-limited by its cognate cell membrane transporters (encoded by SLC2A gene family). RNA sequencing (RNA-seq) analysis of gene expression revealed that the most prominently upregulated transporter commonly in resistant cell line models was GLUT6 (encoded by SLC2A6 gene) (Figure 2A), a non-canonical class III glucose transporter rather than the canonical GLUT1 (SLC2A1) or GLUT3 (SLC2A3) ^5, 18, 19^. The GLUT6 mRNA upregulation selectively in resistant cells was validated by reverse transcription quantitative polymerase chain reaction (RT-qPCR) analysis whereas the mRNA expression abundance of the canonical transporters GLUT1 and GLUT3 in general did not show markable difference between the resistant versus parental cells (Figure 2B). Western blot analysis reconfirmed the upregulation of GLUT6 protein in the resistant cells (Figure 2C). Immunofluorescence staining analysis orthogonally demonstrated the upregulation of cell membrane GLUT6 protein location and abundance in resistance cells (Figure S3A). The upregulation was recapitulated in KRAS inhibitor (KRASi)-resistant and tolerant cells (Figure S3B, S3C).

**Figure 2.**
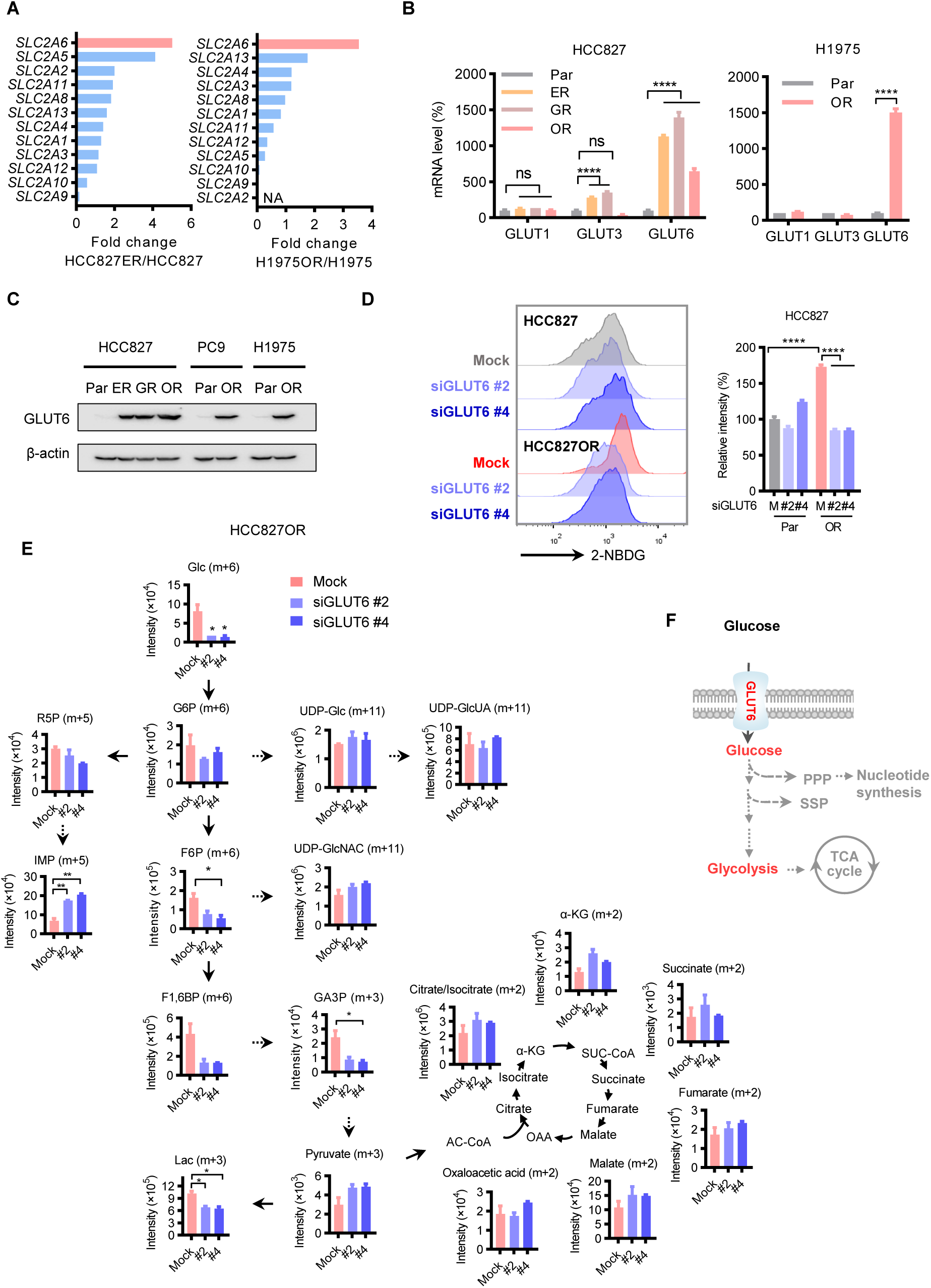
The rewired glucose uptake and metabolism in resistant cells are determined by a non-canonical transporter GLUT6. (A and B) Differentially expressed glucose transporter-encoding genes in resistant versus parental cells assayed by RNA-seq (A) and RT-qPCR (B) analyses. (C) Western blot analysis of GLUT6 protein expression. β-actin used as a loading control. (D and E) Abrogation of enhanced glucose uptake and glycolysis in resistant cells by knockdown of GLUT6 assayed in fluorescent glucose analog 2-NBDG-based flow cytometry (D) and SIRM (E) analyses. Cells were transfected with 20 nM siGLUT6 or Mock control for 48 h. (F) Schematic showing enhanced glucose uptake in EGFRi-resistant cells funneled into the glycolysis pathway determined by GLUT6. For (B), (D), and (E), data are representative of 4, 2, and 2 independent experiments with 3, 4, and 2 technical replicates, respectively, and presented as means ± SEM. *, *P* < 0.05; **, *P* < 0.01; ***, *P* < 0.001; ****, *P* < 0.0001.

Knockdown of GLUT6 with small interfering RNA (siRNA) abrogated the enhanced glucose uptake and glycolysis selectively in resistant cells assayed by 2-NBDG-based flow cytometry analysis (Figure 2D, S3D). This abrogation effect of GLUT6 knockdown on enhanced glucose uptake and glycolysis selectively in the resistant cells was validated in SIRM analysis (Figure 2E) and reconfirmed in Cell Energy Phenotype Test analysis (Figure S3E).

These results indicate that in resistant cells the non-canonical transporter GLUT6 is upregulated and causes glucose uptake enhancement and metabolism rewiring (Figure 2F).

### The non-canonical transporter GLUT6 determines resistance to targeted therapy

Since the rewired glucose uptake and metabolism in resistant cells are determined by GLUT6, we then investigated whether the resistance phenotype is dictated by GLUT6. Knockdown of GLUT6 selectively inhibited EGFRi-resistant cells, showing more inhibition of cell growth (Figure 3A, S3F) and colony formation (Figure 3B) in resistant cells than their parental counterparts. This selective effect of GLUT6 was mirrored in KRASi-resistant cells (Figure 3C). Moreover, GLUT6 knockdown imposed negligible impact on the viability of the normal airway epithelial cells (Figure S4A). On the contrary, knockdown of canonical glucose transporters GLUT1 (Figure S4B left panel) or GLUT3 (Figure S4B right panel) rendered cytotoxicity to normal airway epithelial cells and generally failed to elicited selective inhibition on the resistant cells (Figure S4C, S4D). These results indicate a selective dependence of resistance on GLUT6.

**Figure 3.**
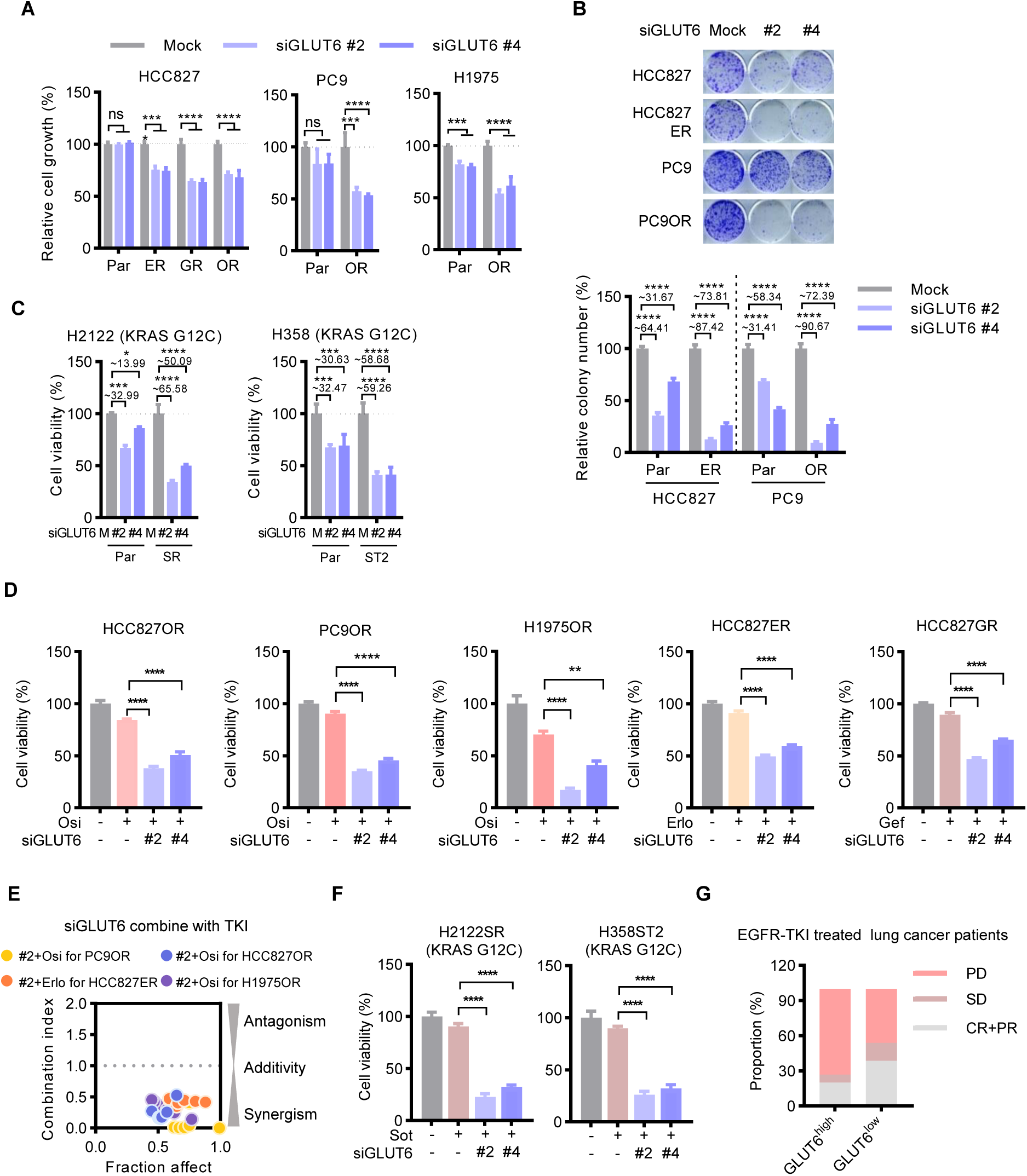
The non-canonical transporter GLUT6 determines resistance to targeted therapy. (A and B) Selective inhibitory effect of GLUT6 knockdown on resistant cells versus parental cells in cell growth and colony formation analysis. Cells were transfected with 20 nM siGLUT6 or Mock control. In grow analysis, the cells were monitored by IncuCyte ZOOM system every 4 hours and the data at 72 h are shown; In colony formation analysis, the treated cells were resuspended and plated for colony formation in drug-free medium for 10 days. (C) More vulnerable to knockdown of GLUT6 in KRASi sotorasib-resistant H2122SR and -tolerant H358ST2 cells versus their counterpart parental cells. SR, sotorasib resistant; ST, sotorasib tolerant. The cells were treated with 20 nM GLUT6 or mock siRNA treatment for 72 h for cell viability analysis based on CCK8 assay. (D) Resensitization of GLUT6 knockdown on EGFR inhibitors in resistant cells. Cell viability assay were performed after cells were treated for 72 h. Osimertinib (Osi), gefitinib (Gef), or erlotinib (Erlo) at 500 nM and GLUT6 or mock siRNA at 20 nM were used. (E) Synergy effect of GLUT6 knockdown on sensitivity to various EGFR inhibitors in HCC827OR, HCC827GR, PC9OR and H1975OR cells. CI value (combination index) was calculated as indicated in the Methods section. (F) Resensitization of GLUT6 knockdown on KRASi sotorasib in resistant and tolerant cells. Cell viability assay were performed after cells were treated for 72 h. Sotorasib at 1 µM in H2122SR (sotorasib resistant) and at 200 nM in H358ST2 (sotorasib tolerant) cells. GLUT6 or mock siRNA at 20 nM. (G) Association of higher expression of GLUT6 with inferior drug response in lung cancer patients underwent EGFRi treatment. Data were retrieved from TCGA dataset (https://portal.gdc.cancer.gov/). The patients were stratified in accordance with high (n=15) versus low (n=13) expression (cutoff: median) of GLUT6 mRNA abundance within their tumors. CR, complete response. PR, partial response. SD, stable disease. PD, progressive disease. For (A), (C), (D) and (F), data are representative of at least 3 independent experiments with at least 3 technical replicates and presented as means ± SEM. *, *P* < 0.05; **, *P* < 0.01; ***, *P* < 0.001; ****, *P* < 0.0001.

We next examined whether GLUT6 is necessary for the inferior response to targeted therapy in resistant cells. Knockdown of GLUT6 resensitized the resistant HCC827ER, HCC827GR, HCC827OR, PC9OR, and H1975OR cells to the corresponding EGFR inhibitors, erlotinib, gefitinib, and osimertinib (Figure 3D, 3E, S4E, S4F). The calculated Combination Indexes (CIs) for the inhibition effect of the combination of GLUT6 knockdown and each inhibitor were less than 1.0 (Figure 3E, S4F), indicating synergy effects ^20^. The resensitization effects of GLUT6 knockdown were recapitulated in KRAS-targeted therapy, where knockdown of GLUT6 resensitized the resistant H2122SR and H358ST2 cells to sotorasib (Figure 3F, S4G, S4H). Consistently, analysis of lung cancer patients underwent EGFRi treatment recorded in The Cancer Genome Atlas (TCGA) database showed that higher expression of GLUT6 in tumors was associated with an inferior drug therapy response (Figure 3G).

### GLUT6-facilitated methylglyoxal derived from glucose metabolism induces resistance

The enhancement of glycolysis was accompanied by an accumulation of lactate in resistant cells (Figure 1F, S2B, S2C), which is determined by GLUT6 as shown by the abrogation of the abundance increases of glycolysis and lactate with GLUT6 knockdown (Figure 2D, 2E, S3D, S3E). Supplementation of lactate in culture media, however, failed to impart resistance in various sensitive cells in viability analysis (Figure 4A, S5A). Coincidently, RNA-seq analysis demonstrated neither the mRNAs of the SLC16A gene family members including the lactate transporters monocarboxylate transporters MCT1 (SLC16A1), MCT2 (SLC16A7), MCT3 (SLC2A8), MCT4 (SLC16A3) nor the lactate dehydrogenases LDHA, LDHB, LDHC, and LDHD were upregulated in various resistant models (Figure 4B). These data imply lactate as a glycolytic epiphenomenon and accompanying biomarker in glycolysis enhancement rather than as a causal factor for targeted therapy resistance.

**Figure 4.**
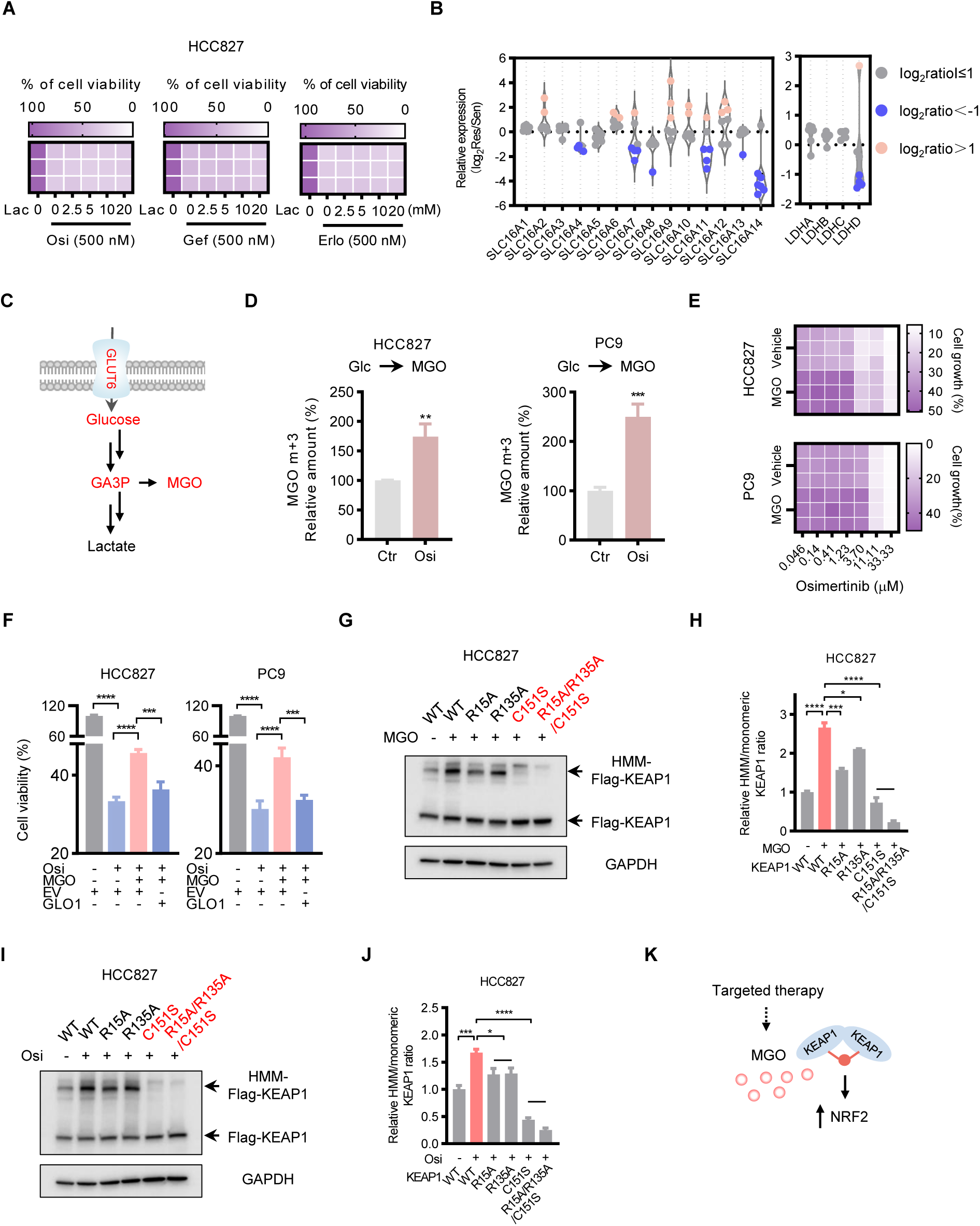
GLUT6-facilitated methylglyoxal derived from glucose metabolism induces resistance and KEAP1 dimerization. (A) Negligible effect of lactate on the sensitivity to EGFR inhibitors in cell viability assay. Cells were treated by lactate (Lac) at indicated concentrations and individual EGFR inhibitors at 500 nM for 72 h. Osi, osimertinib; Gef, gefitinib; Erlo, erlotinib. (B) Transcriptional expression of lactate transporters and lactate dehydrogenases in resistant (HCC827ER, HCC827GR, H1975OR1-5) versus corresponding parental cells assayed by RNA-seq analysis. The definition and description of the resistant cell lines are shown in this article and were indicated in our previously report ^14, 16, 17^. (C) Schematic illustrating glucose metabolism shunt to MGO via GA3P besides conversion to lactate, the well-known pathway. (D) LC/MS-MS analysis of intracellular MGO derived from extracellular medium glucose (MGO m+3). HCC827 and PC9 cells were cultured in the medium where glucose was replaced with the same concentration (11 mM) of U- ^13^C-glucose containing 1 μM osimertinib or solvent control for 6 h and then the intracellular labeled MGO were measured. (E and F) Desensitization effect of MGO on osimertinib response in HCC827 and PC9 cells assayed by growth (E) and viability (F) analysis. After 0.625 mM MGO or vehicle control treatment for 8 h, the cells were resuspended, plated, and exposed to indicated concentrations of (E) or 500 nM (F) osimertinib for 72 h. GLO1, glyoxalase 1 ectopic expression; EV, empty vector. (G and H) Western blot analysis (G) and relative abundance (H) of high molecular mass (HMM) KEAP1 dimer formation of wild-type (WT) or mutant Flag-KEAP1 and its dependent amino acid(s) in HCC827 cells treated with 1.25 mM MGO or solvent control for 8 h. (I and J) Western blot analysis (I) and relative abundance (J) of HMM KEAP1 dimer formation and its determinant amino acid(s) in HCC827 cells exposed to 500 nM osimertinib or solvent control for 36 h. (K) Schematic depicting the hypothesis that the therapy-induced, glucose-derived MGO promotes KEAP1 dimerization, causing an increase of NRF2 protein abundance. For (F), (H) and (J), data are representative of two, two, and three, respectively, independent experiments and presented as means ± SEM. *, *P* < 0.05; **, *P* < 0.01; ***, *P* < 0.001; ****, *P* < 0.0001.

Besides conversion to pyruvate and lactate, glucose via glyceraldehyde-3-phosphate (GA3P) is alternately shunted to the formation of methylglyoxal (MGO) (Figure 4C), a reactive carbonyl species (RCS) compound tending to induce post-translational modifications of proteins ^21–23^. Though not as well-known as lactate-forming metabolism in glycolysis and not a target metabolite considered in usual metabolomic detections, this pathway has been reported to be highly augmented in hyperglycemia and play important roles in diabetes ^21, 23^. We noticed that GA3P stood out in the accumulated intermediate metabolites in glucose metabolism pathways in the established resistant cells (Figure 1F) and in the sensitive cells challenged by drug acute treatment (Figure 1G). This elevation of GA3P in resistant cells was determined by GLUT6 as shown by the abrogation of the elevation after the knockdown of GLUT6 (Figure 1F, 2E).

SIRM analysis by using LC-MS/MS, the current state-of-the art technique for the determination of MGO ^21^, revealed an intracellular increase of MGO derived from extracellular glucose when the cells were challenged with osimertinib treatment (Figure 4D). Moreover, when exposed to MGO, the sensitive cells did acquire resistance to osimertinib (Figure 4E, 4F). This desensitization effect of MGO was abrogated (Figure 4F) by ectopic expression of MGO-eliminating enzyme GLO1 (Figure S5B, S5C), validating the pathway dependence.

### GLUT6-glucose-derived methylglyoxal induces resistance by upregulation of NRF2 via dimerization of KEAP1

MGO has been reported to via modification of KEAP1 upregulate the NRF2 (nuclear factor erythroid 2-related factor 2; encoded by NFE2L2 gene) protein ^22^ which has been proved a drug resistance inducer by our and other groups ^24–26^. We hypothesized that the GLUT6-facilitated MGO induces resistance in a KEAP1-NRF2 axis-dependent manner. MGO treatment induced high-molecular mass (HMM) KEAP1 dimer ^22^ formation (Figure 4G, 4H), which was abrogated by a mutation of the key amino acids, C151, R15, and R135 in KEAP1 protein for HMM-formation (Figure 4G, 4H). Osimertinib, coincident with its induction of glucose-derived MGO formation (Figure 4D), promoted KEAP1 dimerization (Figure 4I, 4J) as MGO did (Figure 4G, 4H) and also depended on the amino acids C151, R15, and R135 in KEAP1 protein (Figure 4I, 4J). We then investigated whether in this situation NRF2 is upregulated (Figure 4K).

Consistent with its resistance-inducing (Figure 4E, 4F) and KEAP1-dimerization (Figure 4G, 4H) effects, MGO upregulated NRF2 protein expression in sensitive cells (Figure S6A, S6B left panel), which was abrogated by ectopic expression of MGO-eliminating enzyme GLO1 (Figure S6C). The NRF2 protein-upregulation effect of MGO was recapitulated by EGFRi treatment (Figure S6B right panel) and depended on the HMM-formation key amino acid(s) (Figure S6B). Time-course western blot analysis showed that in response to drug stress the protein upregulation of NRF2 (began at 48 h and 24 h in EGFRi treatment and KRASi treatment, respectively) followed that of GLUT6 (prior to 24 h and 6 h in EGFRi and KRASi, respectively) (Figure S6D). The upregulation of NRF2 (Figure S6E) and GLUT6 (Figure 2C) protein was sustained and fixed up to resistance-acquired phase though the resistant cells are free from drug treatment challenge in this phase, indicating the augmentation of NRF2 and GLUT6 throughout the process of the resistance initiation and maintenance. The upregulated NRF2 is functional as a transcription factor, as evidenced by an augmented expression of its downstream target gene repertoire in resistant cells (Figure S6F left panel, S6G left panel).

The protein upregulation and hyperactivation of NRF2 in EGFRi-and KRASi-resistant cells were upstream determined by GLUT6 as shown by the abrogation of the upregulated NRF2 protein (Figure S6H) and the augmented NRF2 target gene repertoire expression (Figure S6F right panel, S6G right panel) with GLUT6 knockdown and by the precedent appearance of drug stress-induced protein elevation of GLUT6 ahead of NRF2 elevation (Figure S6D). Accordingly, GLUT6 conferred resistance via NRF2, because the resensitization effect of GLUT6 knockdown to EGFRi inhibitors in resistant cells was abrogated by the treatment of Ki696 (Figure S6I), a small molecule activator of NRF2 ^27, 28^.

These data indicate that GLUT6-facilitated glucose-derived MGO imparts resistance by upregulation of NRF2 via dimerization of KEAP1.

### GLUT6 is transcriptionally upregulated by drug stress-activated MAZ

To identify upstream mechanism that determines GLUT6 mRNA and protein upregulation in resistant cells, we subjected the common differentially expressed genes (DEGs) of various resistant models versus their isogenic parental cells assayed by RNA-seq analysis to transcription factor (TF) prediction according to chromatin immunoprecipitation enrichment analysis (ChEA) based on encyclopedia of DNA elements (ENCODE) and ReMap libraries (Figure 5A). In addition, potential candidate TFs were deduced according to TF binding sites in the promoter regions of GLUT6 gene based on TF-predicting databases (Figure 5A). MAZ stood up as the sole overlapping TF picked up by two assaying systems (Figure 5B).

**Figure 5.**
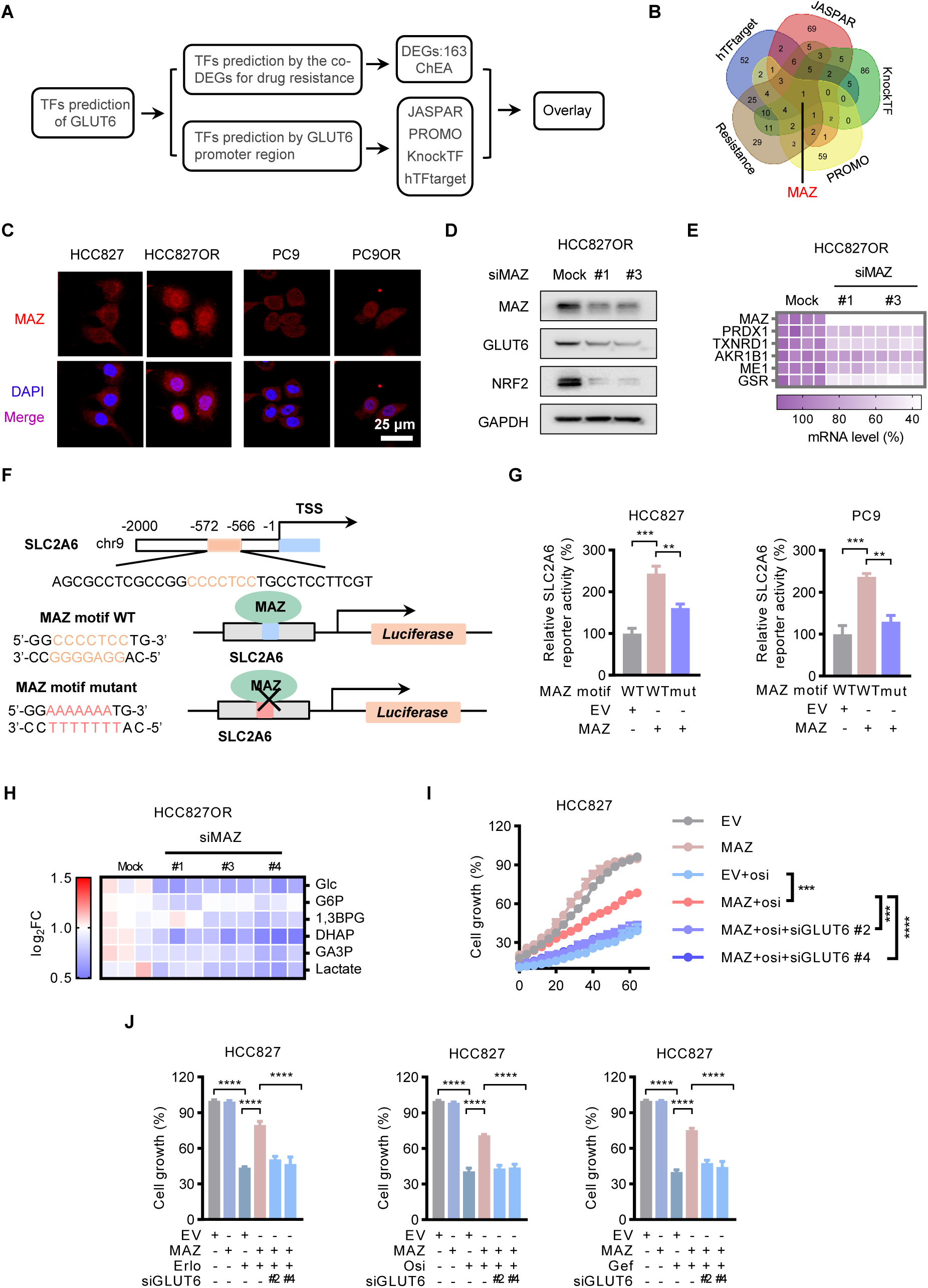
GLUT6 is transcriptionally upregulated by drug stress-activated MAZ. (A) Schematic depicting the screening of transcriptional factors (TFs) that potentially determine GLUT6 upregulation in resistant cells. DEGs, differentially expressed genes; ChEA, chromatin immunoprecipitation enrichment analysis. JASPAR, PROMO, KnockTF, and nTFtarget are databases capable of predicting TF binding to the target gene promoters. (B) Venn diagram shows MAZ as the sole overlapping TF picked up by two assaying systems. (C) Confocal microscopy immunofluorescence staining analysis shows increased nuclear MAZ (red) in resistant compared with parental cells (HCC827OR versus HCC827, PC9OR versus PC9). Nuclei were counterstained with DAPI (4′,6-diamidino-2-phenylindole). (D) Western blot analysis of the regulation effect of MAZ on GLUT6 and NRF2. Cells were transfected with 20 nM MAZ siRNA or mock control for 36 h. (E) Suppression of NRF2 target gene repertoire expression by knockdown of MAZ in resistant cells assayed by RT-qPCR analysis. Cells were transfected with 20 nM MAZ siRNA or mock control for 72 h. (F) Schematic of MAZ-binding sites upstream of transcriptional start sites (TTS) of GLUT6 promoter and the promoter sequences containing the wild-type MAZ-binding motif or the mutated, binding-failed motif cloned with the GLUT6 luciferase reporter. (G) Dual-luciferase reporter analysis of MAZ effect on the promoter activity for GLUT6 in HCC827 and PC9 cells. MAZ, MAZ ectopic expression; EV, empty vector. (H) LC-MS/MS analysis of MAZ effect on glycolysis metabolites abundance in HCC827OR cells. Cells were transfected with 20 nM MAZ siRNA or Mock control for 48 h. (I and J) MAZ ectopic expression-conferred resistance to EGFR inhibitors and the abrogation of GLUT6 knockdown on this effect. Cells were treated with corresponding individual inhibitors at 500 nM, GLUT6 siRNA at 20 nM and monitored and quantified by IncuCyte ZOOM analysis system every four hours for three days (I) or at 72 h (J). Data are representative of three independent experiments with three technical replicates and presented as means ± SEM. *, *P* < 0.05; **, *P* < 0.01; ***, *P* < 0.001; ****, *P* < 0.0001. For (I), statistical significance was assessed using repeated measures two-way ANOVA.

Nuclear translocation is a prerequisite for the functional execution of transcription factors. In situ cellular immunofluorescence staining analysis demonstrated more nuclear MAZ in resistant cells than that in parental cells (Figure 5C). The nuclear translocation of MAZ was initiated by drug stress as shown by an increase of nuclear MAZ abundance in sensitive cells insulted by osimertinib acute treatment (Figure S7A). The effect that MAZ stands upstream controlling GLUT6-NRF2 axis in resistant cells was validated in MAZ knockdown assays where MAZ siRNAs downregulated GLUT6 mRNA expression (Figure S7B), GLUT6 and NRF2 protein expression (Figure 5D), and NRF2 target gene expression (Figure 5E) in these cells.

Dual-luciferase reporter analysis demonstrated that MAZ ectopic expression in sensitive cells increased the luciferase activity driven by GLUT6 (SLC2A6) promoter whereas this effect was lost after a mutation of the promoter MAZ-binding motif (Figure 5F, 5G), validating the direct transcriptional regulation of GLUT6 gene expression by MAZ. In line with the MAZ dictation on GLUT6, knockdown of MAZ suppressed glycolysis (Figure 5H), colony formation (Figure S7C), viability (Figure S7D), and restored sensitivity to osimertinib (Figure S7E) in resistant cells. In contrast, ectopic expression of MAZ in parental cells (Figure S7F, S7G) induced the otherwise sensitive cells to be resistant to the EGFRi treatment (Figure 5I, 5J). This resistance induction effect of MAZ was abrogated by knockdown of GLUT6 (Figure 5I, 5J), confirming its dependence on GLUT6 for resistance.

### Drug resistance acquisition and maintenance in vivo depend on GLUT6

We then investigated the role of GLUT6 in acquisition and maintenance of resistance in vivo by conditionally interfering with GLUT6 (Figure 6A).

**Figure 6.**
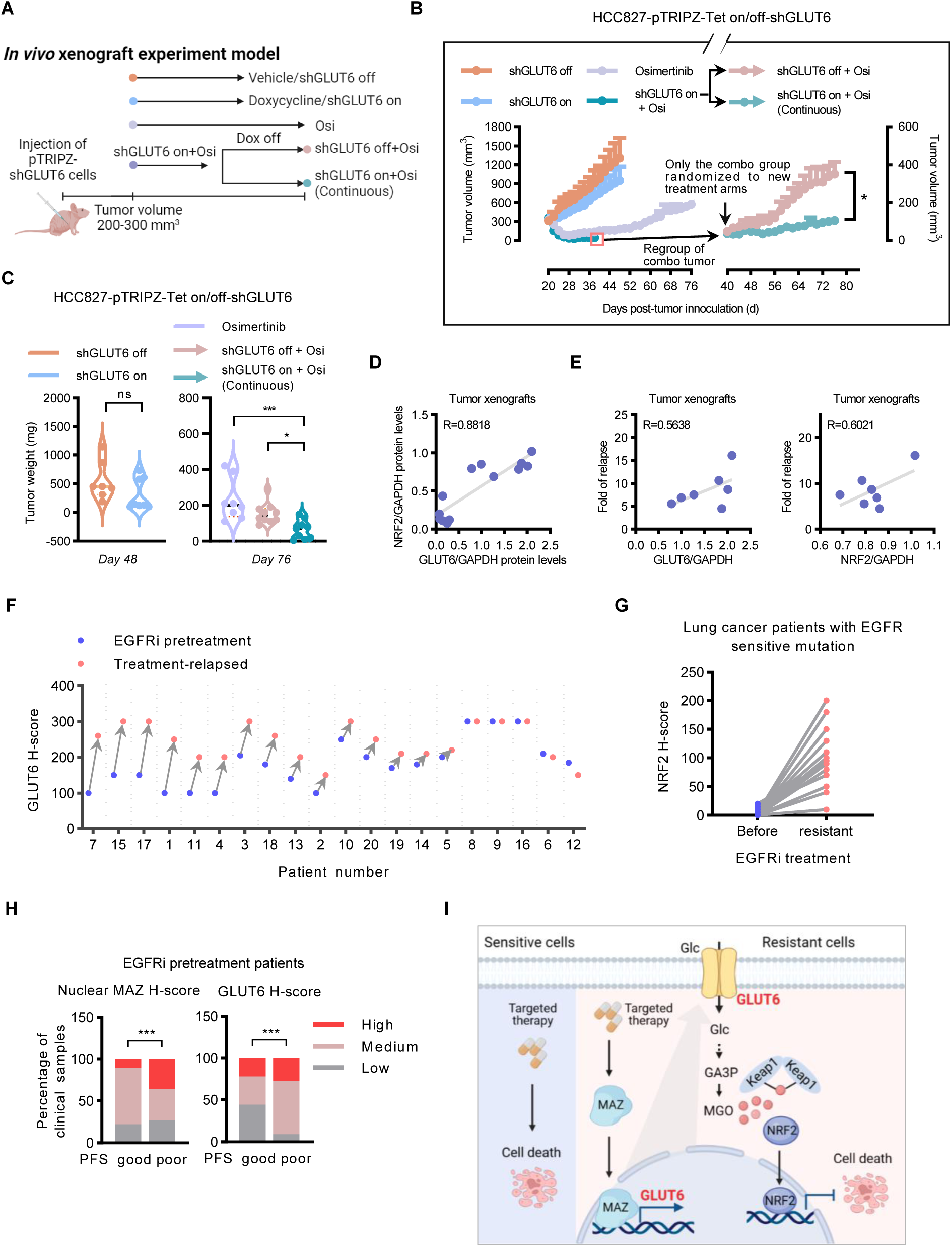
Drug resistance acquisition and maintenance in vivo depend on GLUT6. (A) Schematic overview of the effect of conditional interference of GLUT6 expression on resistance acquisition and maintenance. HCC827 cells expressing doxycycline (Dox) inducible Tet-on-shRNAs, where GLUT6 can be silenced by Dox-induced shRNA expression, were subcutaneously inoculated into the flanks of nude mice (two tumors per mouse). When the tumors reached 200-300 mm^3^, animals were randomized into groups as indicated. (B and C) Effect of conditional knockdown of GLUT6 with doxycycline-inducible shRNAs (Tet-on-shGLUT6) on prevention of osimertinib resistance in HCC827 cell-derived xenograft (CDX) tumors shown as tumor growth monitoring (B) and tumor weight (C) analyses. Groups: shGLUT6 off (GLUT6 non-knockdown administered with vehicle control, n=5 animals); shGLUT6 on (GLUT6 knockdown induced by Dox, n=5 animals); Osimertinib (n=5 animals); and shGLUT6 on + Osi (GLUT6 shRNA + osimertinib, n=10 animals). At day 40 post tumor inoculation, 10 mice in “shGLUT6 on + Osi” group were randomly split and regrouped into shGLUT6 on (continuous) + Osi arm (n=5) where the knockdown was continued with Dox administration and shGLUT6 off + Osi arm (n=5) where the knockdown was turned off by Dox withdrawal. Vehicle (1% dimethyl sulfoxide, once daily), doxycycline (50 mg/kg, once daily), or osimertinib (2 mg/kg, once daily) was administered through oral gavage. Statistical differences were conducted with repeated measures one-way ANOVA (B) and ANOVA followed by Tukey posttest comparison (C). (D) Positive correlation of NRF2 with GLUT6 protein abundances measured by western blot analysis of tumor samples in (B) and (C). (E) Positive correlation of relapse magnitude with GLUT6 (left panel) or NRF2 (right panel) protein abundance in Osi group (tumors shown in (B)). Relapse magnitude is represented as (Vterm-Vmin)/Vmin. Vterm, the tumor volume in relapse and termination of the test; Vmin, the tumor volume in maximal response to osimertinib treatment. (F and G) Evaluation of protein abundance of GLUT6 (F) and NRF2 (G) in the matched EGFRi pretreatment and treatment-relapsed tumor sample pairs in the same patient. Forty paired specimens from EGFR-mutant lung cancer patients treated with EGFRi were used for immunohistochemistry (IHC) staining. Protein expression was semi-quantified in a blind manner by the pathologist presented as the histoscore (H-score), ranging from 0 to 300, calculated based on the staining intensity of target molecule(s) and the percentage of stained tumor proportion. (H) High expression of cellular GLUT6 and nuclear MAZ in pretreatment tumors predicted worse treatment response. PFS, progression-free survival. Poor response, PFS < 16 month; good response, PFS > 16 month; (cutoff: medium). Twenty patients with EGFR-mutant lung cancers underwent EGFR TKI targeted therapy were analyzed. (I) Schematic demonstration of the drug resistance conferred by therapy stress-MAZ-GLUT6-glucose noncanonically metabolic rewiring-MGO-KEAP1-NRF2 axis. *, *P* < 0.05; **, *P* < 0.01; ***, *P* < 0.001; ****, *P* < 0.0001.

HCC827 cell-derived xenograft (CDX) tumors showed an initial response to osimertinib with tumor volume shrink and grow delay but then acquired resistance as demonstrate by a rebound of tumor burden during the continuous treatment (Figure 6B, “Osimertinib” cohort). Doxycycline (Dox)-inducible knockdown of GLUT6 (Figure S8A) blocked the tumor relapse on osimertinib (Figure 6B and 6C, “shGLUT6 on + Osi” cohort and “→ shGLUT6 on (Continuous) + Osi” cohort versus “Osimertinib” cohort) though the knockdown itself without the combination of osimertinib treatment only had negligible effect on tumor growth (Figure 6B and 6C, “shGLUT6 on” cohort vs. “shGLUT6 off” cohort), indicating the dependence of drug resistance acquisition on GLUT6. This effect was reconfirmed when the knockdown was turned off by withdrawal of doxycycline, where the blocked rebound was unleashed as shown by a regrowth of the suppressed tumor burden (Figure 6B and 6C, “→ shGLUT6 off + Osi” Arm vs. “→ shGLUT6 on (continuous) + Osi” Arm). Western blot analysis of osimertinib treatment-relapsed tumors demonstrated an increase of GLUT6 and NRF2 protein abundance (Figure S8B), mirroring the in vitro results where the cancer cells upregulated GLUT6 and NRF2 protein expression during the drug resistance acquisition (Figure 2C, S6D, S6E). In tumors, NRF2 protein abundance showed a positive correlation with that of GLUT6 (Figure 6D). In osimertinib-treated tumors, replase magnitude positively correlated with the protein abundance of GLUT6 or NRF2 (Figure 6E). These data indicate that GLUT6 is necessary for drug resistance and as a potential target for resistance prevention.

We next sought to examine whether the resistance maintenance is determined by GLUT6 and whether targeting GLUT6 overcomes the resistance. In the resistance-established HCC827OR cell-derived xenograft (HCC827ORCDX) tumors osimertinib as a single agent failed to shrink the tumor volume (Figure S8C) as in the HCC827CDX (Figure 6B), however, when combined with GLUT6 knockdown (Figure S8E), fulfilled complete shrink and continued repression of tumor growth rebound (Figure S8C, S8D). Importantly, when the knockdown was turned off (Figure S8E), the resistance recurred (Figure S8C, S8D). These data indicate the dependence on GLUT6 of the resistance maintenance and imply a drug resistance-overcoming promise of targeting GLUT6.

To investigate the clinical relevance of the augmented MAZ-GLUT6-NRF2 signaling in therapy resistance, we examined these signaling molecules in matched pairs of pretreated and EGFRi treatment-relapsed non-small-cell lung cancer (NSCLC) specimens. Immunohistochemistry (IHC) analysis showed an upregulated protein expression of tumor cell GLUT6 and NRF2 (Figure 6G, S8F) when the patients relapsed to the treatment. Moreover, higher expression of GLUT6 and nuclear MAZ in pretreatment tumors was associated with worse progression-free survival after treatment (Figure 6H, S8G), indicating its predictive implications for drug response and relapse.

## Discussion

Targeting glucose metabolism for cancer therapy is a long-standing quest though stumbles largely due to lacking of enough understanding of the distinctions between the malignancy traits to enable selective intervention for therapeutic margin. We find that when cancer cells acquire resistance to targeted therapy, they coincidentally acquire addiction to a non-canonical glucose transporter GLUT6 for glucose uptake and utilization. The resistant cells upregulate and take advantage of GLUT6 to counteract drug stress while expose this metabolic vulnerability that can be exploited and leveraged by us to selectively attack these resistant cells.

Glucose transporters, as the pivot for glucose entry into the cells and as the cell membrane proteins available for extracellular molecule binding, have long been the enthusiasm in research and therapy exploitation. The canonical class I GLUTs, such as GLUT1 and GLUT3 have received more intensive investigation for cancer interception ^29–31^, though they are ubiquitously expressed across tumor and non-tumor cells, important for general glucose homeostasis, and less possible to fulfill a therapeutic margin ^19^. GLUT6, as a class III glucose transporter, selectively upregulated in and required by resistant cell revealed in this work, has been reported low expressed and not a major regulator for general body metabolism in physiological conditions ^32, 33^, thus showing opportunity as an ideal specific target for drug resistance overcoming.

We found that GLUT6 conferred resistance by facilitating glucose influx and conversion to methylglyoxal which bond to KEAP1 to relieve the degradation induction effect of KEAP1 on NRF2. Imported by GLUT6, the glucose with increased abundance was converted to more methylglyoxal. This is mirrored by the discovery of methylglyoxal formation augmentation in diabetic hyperglycemia conditions ^21, 34^. The resultant NRF2 pathway activation protected the cancer cell from drug insults and conferred resistance, reminiscent of our and other group findings where NRF2 drives drug resistance through transcription of cytoprotective genes ^24–26^.

Coincided with the above-mentioned downstream mechanism, we found that GLUT6 is upstream transcriptionally upregulated by MAZ which is activated by drug insult. Glucose transporters are predisposed to be regulated to adapt to various conditions. GLUT1, the major glucose transporter ubiquitously expressed in tumor and non-tumor cells at steady state without drug challenge, is determined by HIF-1 and NF-κB transcriptional regulation ^35, 36^ while GLUT3, the dominate isoform in brain cancer cells, transcriptionally upregulated by YAP/TAZ in glioblastoma ^37^. MAZ is a stress- and inflammation-induced zinc finger transcription factor whose physiological and pathological functions have not been well studied ^38, 39^.

Together, we identify that targeted therapy stress-induced activation of MAZ transcriptionally upregulates GLUT6 and that GLUT6 facilitates glucose influx and subsequent increase formation of methylglyoxal which boosts KEAP1 dimerization and activates NRF2 pathway to confer resistance. The cancer cells that utilize this mechanism acquire survival advantage and resistance to the therapy but at the expense of getting addiction to this metabolic reprogramming that can be exploited and leveraged by us to selectively attack the resistance in a metabolic vulnerability-targeted manner (Figure 6I). This finding is of significance. First, we find an unrecognized, nongenetic, metabolic rewiring driving cancer resistance to targeted therapy and have dissected the underling mechanism. This reveals a new dimension of mechanism, other than the well-recognized genetic alterations, for molecularly targeted therapy resistance, the important and urgent clinical and cancer biology question. Second, we find a non-canonical transporter GLUT6-rewired glucose metabolism and shunt to methylglyoxal beyond the well-known canonical transporters and canonical pyruvate-lactate flow. In addition, we have delineated their running scenario —— selectively required and sufficient for resistance to cancer targeted therapy. The underlying upstream and downstream mechanism has also been dissected by us. Last, this discovery that the preferential dependance of resistant cancer on the non-canonical, general-homeostasis-less-perturbing transporter GLUT6 implies a resistance-selective target capable of potentially achieving therapeutic margin and promises a chance for glucose metabolism targeting strategy, a long quest in in cancer therapy and biology science.

## Materials and Methods

### Cells and cell culture

Human lung cancer cell lines harboring EGFRi-targetable mutation(s), HCC827 and H1975 were obtained from the American Type Culture Collection (ATCC), PC9 from Dr. Q. Dong (China State Key Laboratory of Oncogenes and Related Genes). The KRAS-G12C mutated human H2122 and H358 cell lines were from Zhong Qiao Xin Zhou Biotechnology and Cell Bank of the Chinese Academy of Sciences (Shanghai, China), respectively. Human normal lung epithelial cell line BEAS-2B was from FuHeng Cell Center (Shanghai, China). The cells were authenticated by short tandem repeat (STR) analysis. The isogenic resistant cell lines insensitive to the first-generation EGFRi gefitinib (HCC827GR), erlotinib (HCC827ER), and the third-generation TKI osimertinib (HCC827OR, PC9OR, and H1975OR) were established, maintained, and authenticated as indicated (Figure S1D, S1E) and previously described ^14, 16, 17^. H2122SR is the H2122-derived cell line acquired resistance to sotorasib; H358ST2 is the line tolerant to 2000 nM sotorasib; resistance and tolerance are defined and described as in our previous article ^17^.The cells were cultured at 37°C with 5% CO_2_. HCC827, PC9, H1975 and their isogenic resistant cells were cultured in RPMI 1640 medium. BEAS-2B cells were in DMEM medium. The media were supplemented with 10% (vol/vol) fetal bovine serum (FBS), 1% penicillin and streptomycin, and 1% GlutaMax.

### Clinical information

Positron emission tomography-computer tomography (PET-CT) information at relapse phase versus paired non-replase (response and pre-treatment) phase, paired EGFRi pretreatment and treatment-relapsed tumor samples for immunohistochemistry (IHC) staining analysis, and patient information for progression-free survival (PFS) analysis were from the Chest Hospital Affiliated to Shanghai Jiao Tong University School of Medicine and approved by the clinical Ethics Committee. PET-CT measure of glucose uptake and utilization is calibrated with analysis of the maximum standardized uptake value (SUVmax) and quantification of IHC specimens are scored with H-score described in corresponding figure legend section.

### Animal study and in vivo assay

Animal studies were approved by the Institutional Animal Care and Use Committee (IACUC) and performed in compliance with the institutional guidelines of Shanghai Jiao Tong University School of Medicine. For xenograft formation, cells were subcutaneously inoculated into the flanks of 5-week-old BALB/c nu/nu athymic mice. Tumors were detected with a caliper every 2 or 3 days to calculate the volumes according to the formula volume = length × width^2^ × 0.5. When tumor volume reached 200-300 mm^3^, mice were randomly allocated to various treatment groups as described in corresponding figure legend section.

### Cell viability and growth assay

Cells were seeded in 96-well plates. Cell viability were analyzed by using the Cell Counting Kit-8 (CCK-8, Dojindo) in accordance with the manufacturer’s guidelines. Cell growth was monitored and quantified by using the IncuCyte ZOOM live cell system (Essen Bioscience) every 4 h.

### Colony formation assay

Cells dissociated into single-cell suspensions were seeded at 1000 per well in six-well plates for incubation for 7-10 days. The cells were fixed with 4% paraformaldehyde and stained with crystal violet for counting with ECLIPSE TI-S imaging system (Nikon).

### RNA extraction and quantitative polymerase chain reaction (qPCR)

Total RNA from cells was extracted using the RNA extraction kit (Takara) according to the manufacturer’s instructions. Reverse transcription was performed by using the PrimeScrip RT Master Mix Kit. Real-time PCR were carried out using SYBR Premix Ex Taq (Takara) and run in LightCycler 480 II system (Roche). The primer sequences are shown in Table S1.

### Western blot analysis

Cell and tumor samples were lysed on an ice by using radioimmunoprecipitation assay (RIPA) buffer (Beyotime) supplemented with protease and phosphatase inhibitors. Protein abundance was measured with bicinchoninic acid (BCA) assay kit (Thermo Fisher). The proteins were separated by SDS–polyacrylamide gel electrophoresis (SDS-PAGE) and transferred to polyvinylidene difluoride membranes. The membranes were blocked in 5% non-fat milk for 1 h at room temperature and incubated with primary antibody overnight at 4°C. After incubation with corresponding secondary antibodies, the membranes were subjected to electrochemiluminescence (ECL) and scanned by using the imaging analysis system.

### Gene knockdown by using RNA interference

Cells were seeded in 6-well or 96-well plates at a density of 200000 or 4000 cells/well, respectively. Small interfering RNA (siRNA) diluted in Opti-MEM serum-free medium and incubated with lipofectamine 3000 for 5 min at room temperature was added to the cell culture medium. Forty-eight hours later, silencing efficiency was detected by using real-time qPCR and western blot analysis. The sense sequences of shRNAs are shown in Table S1.

### Immunofluorescence and immunohistochemical staining

Cells grown on glass-bottomed dishes were fixed with 4% paraformaldehyde for 15 min, permeated with 0.4% Triton for 1 h, blocked with 5% BSA for 1h, incubated with anti-MAZ antibody (ImmunoWay) overnight, and incubated with Alexa Fluor 555-labeled donkey anti-rabbit IgG(H+L) secondary antibody for 1h, stained with DAPI (Beyotime) for 15min, and then subjected to laser confocal microscope imaging analysis (Leica TCS SP8). Immunohistochemical staining (IHC) was performed on formalin-fixed paraffin-embedded tissue sections (3.5 µm thickness) from EGFR-mutant lung cancer patients. IHC for GLUT6 (1:50, Sigma Cat# SAB2102200), NRF2 (1:50, abcam Cat# ab62352) and MAZ (1:25, Immunoway Cat# YT2669) was performed using the Ventana BenchMark ULTRA instrument (Ventana/Roche, AZ, USA) following the manufacturer’s instruction. The degree of protein expression in tumor cells was assessed by two experienced pathologists corresponding to the H&E slides based on a histoscore (H-score) method, which was calculated by multiplying the percentage of stained tumor area (1-100%) and the staining intensity score from 0 to 3, ranging from 0 to 300.

### Luciferase reporter assay

Cells were transfected with MAZ and the corresponding empty vector (EV) plasmid with lipofectamine 3000 reagent. Forty-eight hours later, the medium was replaced by fresh medium containing puromycin for the selection of the positive clones. EV and MAZ overexpressed cells were seeded in 24-well plates at density of 50000 cells per well. The next day, co-transfected with SLC2A6 WT or Mutant reporter and pGL4.74 hRluc luciferase reporter (E6921, Promega), the cells were incubated at 37°C for 48 h. Cells were assayed according to the instructions of the manufacturer (Dual-Luciferase® Reporter (DLR™) Assay System, E1910, Promega).

### Glucose uptake assay

Measurement was performed by using the Glucose Uptake Cell-Based Assay Kit (Cayman) according to the manufacturer’s instructions. Following incubation with the fluorescent glucose analog 2-NBDG [2-(N-(7-nitrobenz-2-oxa-1,3-diazol-4-yl) amino)-2-deoxyglucose]) for 30 min, the cells were digested and re-suspended, subjected to flow cytometry analysis.

### Liquid chromatography and mass spectrometry

Metabolites were measured using the ExionLC^TM^ AD UPLC system (SCIEX, USA) coupled to a TripleTOF^TM^ 6600 Plus mass spectrometer (SCIEX, USA) via electrospray ionization with the following source conditions: source temperature, 550°C (ESI+) and 450°C (ESI−); ion spray voltage, 5500 (ESI+) and −4500 V (ESI−); atomization gas pressure, 55 psi; auxiliary heating gas pressure, 55 psi; curtain gas pressure, 35 psi. The m/z range was 70 to 1000. Data were acquired using Analyst^TM^ software (SCIEX, USA). Metabolite identification and quantitation were performed using the SCIEX OS^TM^ 1.0 software (SCIEX, USA). The incorporation of ^13^C from U-^13^C-glucose induces an intensity shift from the unlabeled position (m + 0) to the labeled position (m + n, ‘n’ means the number of incorporated ^13^C atoms). The natural isotope abundance correction was made by the AccuCor R package (https://github.com/XiaoyangSu/ IsotopeNatural-Abundance-Correction).

### Glucose metabolism analysis and glucose metabolism stable isotope-resolved metabolomics (SIRM) analysis

Cells were cultured in glucose-free RPMI 1640 medium supplemented with 10% FBS containing 11 mM glucose (for glucose metabolism) or U-^13^C-glucose (for glucose metabolism SIRM analysis) for 6 h. Metabolites in glucose metabolism were extracted from cells. Placed on ice, cells were washed twice with cold PBS buffer and then 500 µL of ice cold 80% methanol was added to the cell dish. Cells were scraped, and vortexed for 10 min at 4°C, and then centrifuged at 20,000 g for 10 min at 4°C. The supernatants were concentrated in a vacuum centrifugal concentrator for 2-3 h, and resuspended in 200 µL of a 50/50 (v/v) acetonitrile/water mixture. Metabolites were separated by injecting 5 μL of sample on a SeQuant^TM^ ZIC-HILIC Polymeric column (2.1*150 mm, 5µm, EMD Millipore). Mobile phase A consisted of 20 mM ammonium carbonate in water and B was acetonitrile. The gradient program was set in the following: 0-1 min.: hold at 10% A, 1–25 min.: linear gradient from 10% to 40% A; 25.1–33min.: hold at 10% A.

### Glucose-derived methylglyoxal (MGO) analysis

Cells were cultured in glucose-free RPMI 1640 medium supplemented with 10% FBS containing 11 mM U-^13^C-glucose for 6 h. Metabolites from the harvested cells were extracted and centrifuged at 20,000 g for 10 min at 4°C. The supernatants were evaporated to dryness. Two hundred microliter acetonitrile were used to reconstitute the dry extracts and then mixed with 200 μL of 1 mg/mL 2,4-dinitrophenylhydrazine (DNPH) in 50% acetonitrile aqueous solution for MGO derivatization. After DNPH derivatization, the mixtures were evaporated to dryness. The dried derivatized metabolites were dissolved in 200 μL acetonitrile for the following analysis: Metabolites (MGO m+3 et.al) were separated by injecting 5 μL sample on a ZORBAX SB-C18 column (2.1*50 mm, 5 µm, Agilent). The mobile phase was as follows: (A) 0.1% (v/v) formic acid in water and (B) 0.1% (v/v) formic acid in acetonitrile. The gradient program was set in the following: 0-1 min.: hold at 30% B, 1-12 min.: linear gradient from 30% to 100% B; 12.1–15 min.: hold at 30% B.

### Extracellular acidification rate (ECAR) and Cell Energy Phenotype Test analyses

The extracellular acidification rate (ECAR) and Cell Energy Phenotype Test were performed by using seahorse XFe96 analyzer (Agilent Technologies) according to the manufacturer’s kits and instructions. The day prior to assay, cells were seeded in cell plates at density of 20000 cells/well and the sensor cartridge were hydrated in XF calibrant at 37°C in a non-CO_2_ incubator overnight. 24 hours later, cells medium was replaced by assay medium and incubated in non-CO_2_ incubator for 1 hours. Compounds were prepared and loaded into the appropriate ports of hydrated sensor cartridge. For ECAR assays, compounds were injected during the measurements as follows: Glucose (10 mM), oligomycin (1 μM) and 2-DG (50 mM). For energy phenotype assays, oligomycin (1 μM), FCCP (0.125 μM). The ECAR and OCR values were normalized to the number of cells.

### Combination effect analysis

Treatment combinations demonstrate as a synergistic, antagonistic, additive, or potentiative effect. The effects were evaluated by a combination index (CI) calculation in accordance with the Chou-Talalay method ^20^. Data were analyzed by using the CompuSyn software (CompuSyn Inc.): CI = 1, additive effect. CI > 1, antagonistic effect. CI < 1, synergistic effect (0.85 to 0.9, slight synergism; 0.7 to 0.85, moderate synergism; 0.3 to 0.7, synergism; 0.1 to 0.3, strong synergism; CI < 0.1, very strong synergism).

### Statistical assay

Data are demonstrated as means ± SEM analyzed with Prism software (GraphPad). Statistical differences were conducted with the two-tailed Student’s t test comparing two treatments or ANOVA followed by Tukey or Dunnett posttest comparing more than two treatments unless otherwise mentioned. *P* < 0.05 was considered as statistically significant. All correlations were calculated using Pearson correlation.

## Acknowledgments

We thank the support of Shanghai Collaborative Innovation Center for Translational Medicine and L. Zheng (Children’s National Medical Center, Shanghai) for helpful comments.

## Author contributions

H.L. and R.X. established in vitro and in vivo experiment conditions, performed analyses for cell growth, viability, western blot, and in vivo response under drug treatment. H.L. performed analyses for TCGA, RNAseq, and omics data mining. H.L. H. L, and P.Z. cultured cells, performed analyses for cell phenotypes. Y.T. and A. A. performed LC-MS/MS– based chemical analysis and metabolomic analysis. J. L., C. X., J. L., W. Z., B. H., and H. Z. contributed to pathological and clinical information collection, analysis, and interpretation. Y.S. provided study materials and contributed to project discussion. H.L. designed the study, analyzed and assembled data with R. X’s help. H.C. contributed to oversight and leadership responsibility for the research and approved the manuscript. L.Z. designed study conception and experiments, assembled and interpreted data, wrote the manuscript, and approved the manuscript.

## Data and materials availability

All data associated with this study are present in the paper or the Supplementary Materials.

## Supplementary Figures

**Figure S1.**
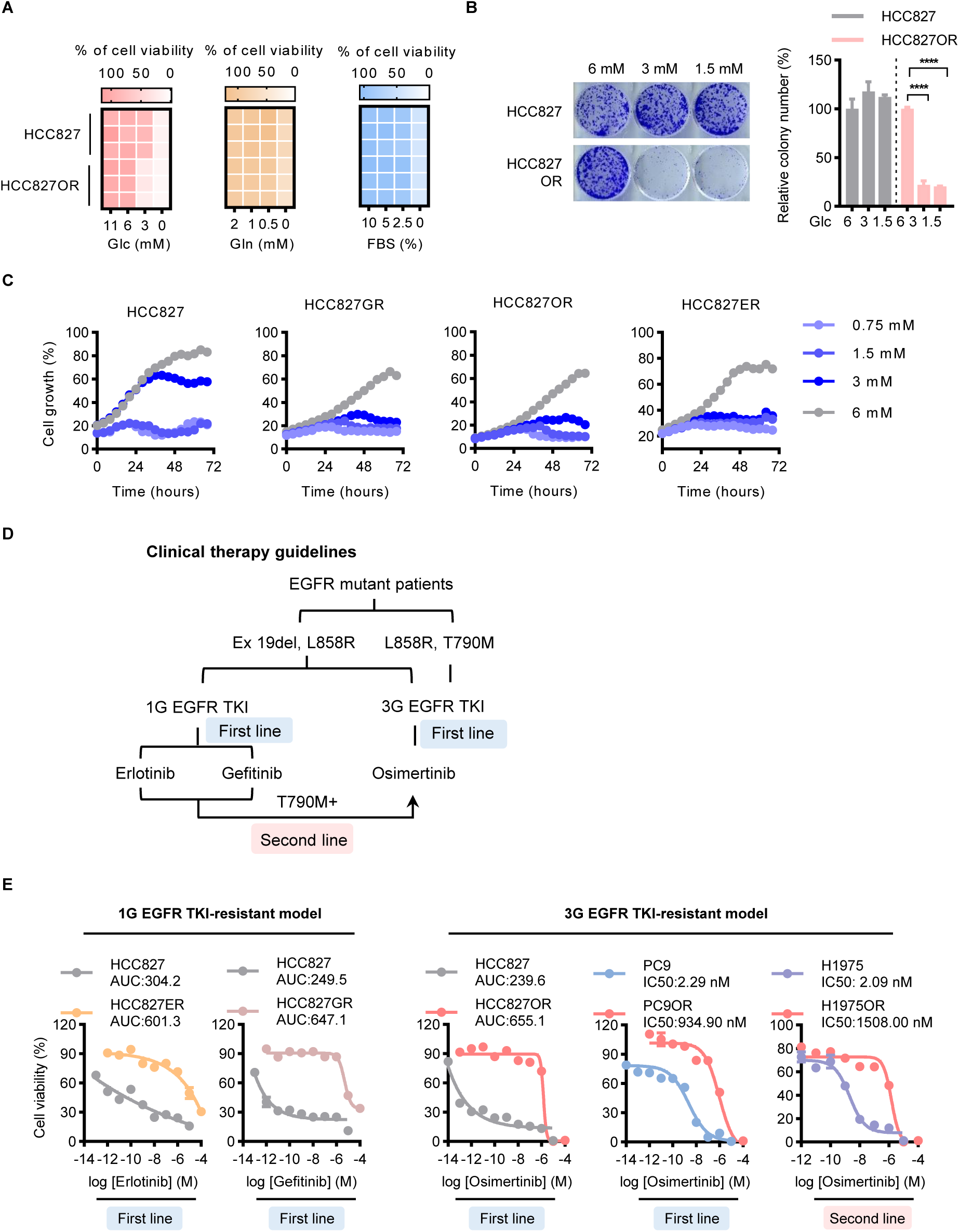
Additionally exacerbated addiction to glucose in cancer resistance to molecularly targeted therapy. (A) Selective vulnerability to decreased medium glucose (Glc) rather than to glutamine (Gln) or fetal bovine serum (FBS) in HCC827OR versus counterpart parental cells in viability analysis. OR, osimertinib resistant. (B and C) More vulnerable to decreased medium glucose in resistant cell survival and growth than in parental cells assayed by colony formation (B) and IncuCyte ZOOM imaging real-time (C) analyses. Colony formation was measured on day 10 after the cells were incubated in medium containing indicated concentrations of glucose. IncuCyte imaging monitored ever 4 h for 72 h. (D) Non-small-cell lung cancer (NSCLC) EGFR-targeted therapy algorithm based on clinical practice guidelines from National Comprehensive Cancer Network (NCCN) and the review article referenced in the main text. TKI, tyrosine kinase inhibitor. (E) Resistance was demonstrated by a right shift of the curve of the TKI concentrations - cell viability inhibition in resistant versus their corresponding parental cells. The cells were treated for 72 h with indicated TKI and concentrations. The cell viability was measured by CCK-8 assay. The half-maximal inhibitory concentration (IC50) or the area under the curve (AUC) of the drug concentration-effect, as a quantitative metric for drug activity ^40, 41^, were properly presented. For (B), data are representative of two independent experiments with two technical replicates and presented as means ± SEM. *, *P* < 0.05; **, *P* < 0.01; ***, *P* < 0.001; ****, *P* < 0.0001.

**Figure S2.**
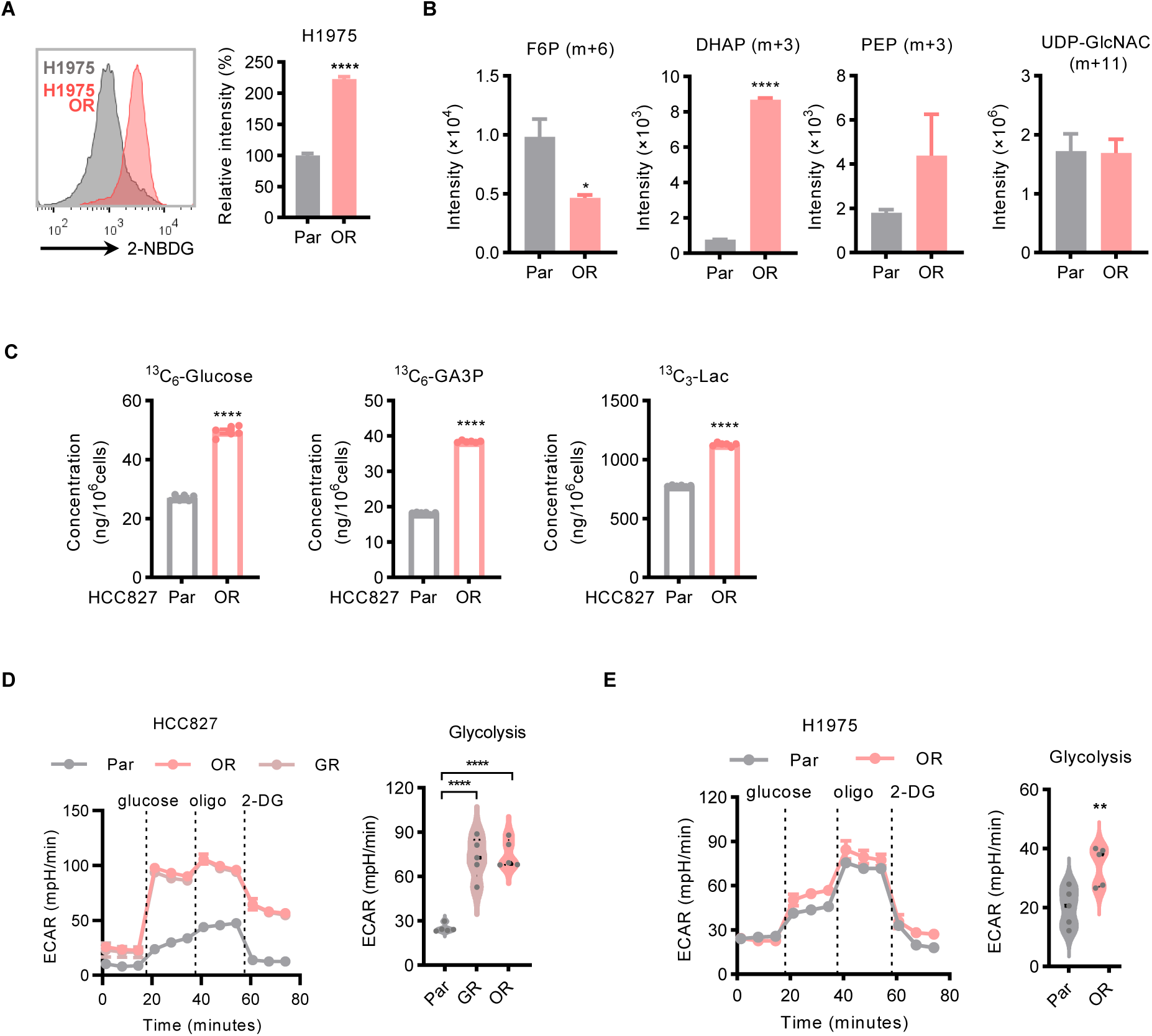
Additional enhancement of glucose uptake and glycolysis in resistant versus sensitive cells. (A) Enhanced intracellular accumulation of medium fluorescent glucose analog 2-(N-(7- nitrobenz-2-oxa-1,3-diazol-4-yl) amino)-2-deoxyglucose (2-NBDG) in H1975OR versus H1975 cells assayed by using flow cytometry. (B) Enhancement of labeled glucose uptaken and funneled into glycolysis in resistant cells versus parental cells. Cells were cultured in medium where glucose was replaced with the same concentration (11 mM) of U-^13^C-glucose for 6 h. The labeled metabolites were determined by LC/MS-MS-based stable isotope–resolved metabolic analysis (SIRM) analysis. (C) Absolute quantification of the increased abundance of ^13^C_6_-glucose, ^13^C_3_- GA3P and ^13^C_3_-Lactate in HCC827OR cells compared with its parental HCC827 cells. Cells were cultured in medium where glucose was replaced with the same concentration (11 mM) of U-^13^C-glucose for 6 h. (D and E) Elevated glycolysis in HCC827 (D) and H1975 (E) resistant versus parental cells assayed by live-cell metabolic analysis of the extracellular acidification rate (ECAR) representing glycolysis by using Seahorse XFe96 analyzer. OR, osimertinib resistant; GR, gefitinib resistant. For (B), (C), (D) and (E), data are representative of at least 3 independent experiments with at least 3 technical replicates and presented as means ± SEM. *, *P* < 0.05; **, *P* < 0.01; ***, *P* < 0.001; ****, *P* < 0.0001.

**Figure S3.**
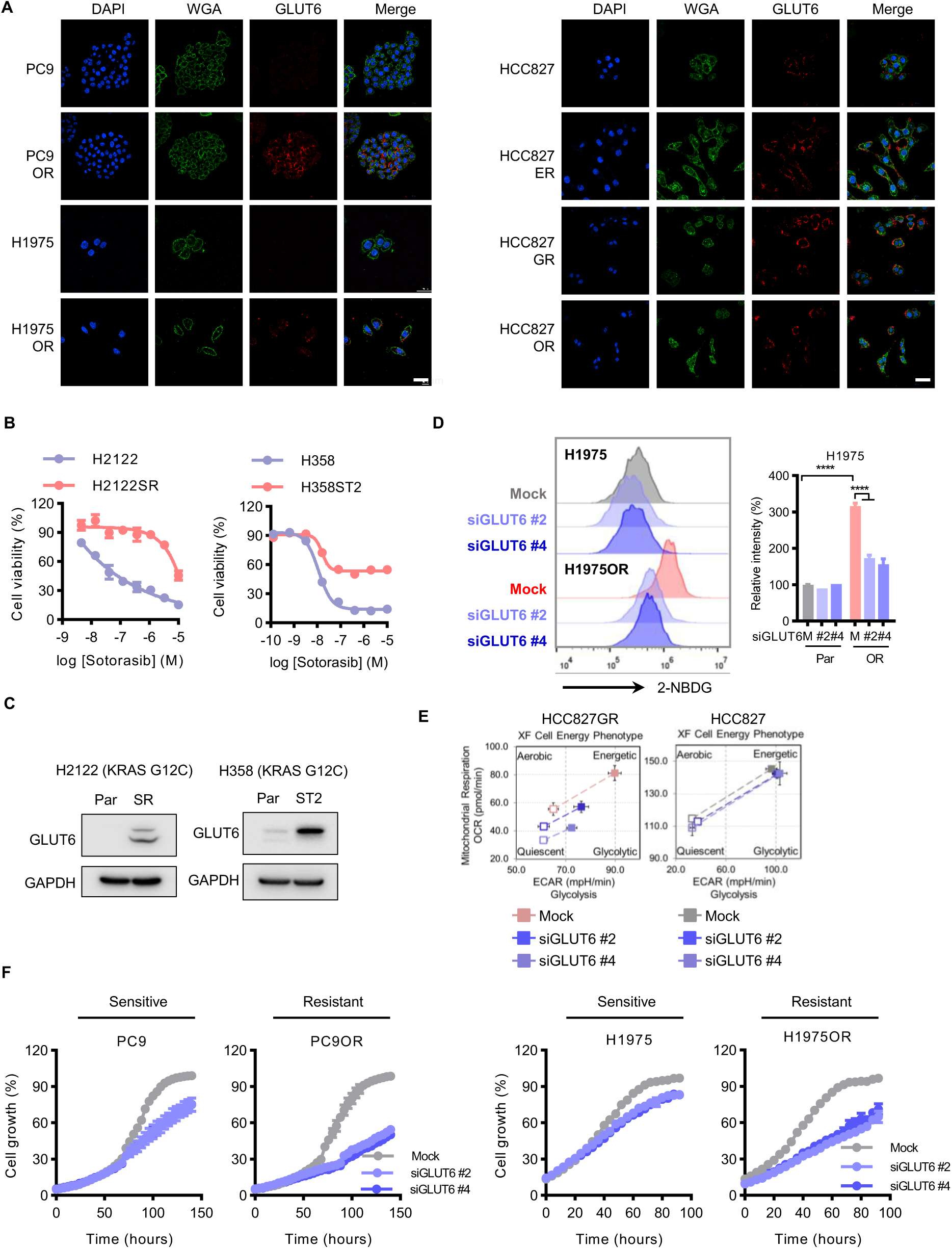
The rewired glucose uptake and metabolism in resistant cells are determined by a non-canonical transporter GLUT6. (A) Upregulation of cell membrane GLUT6 protein (Red) location and abundance in resistant versus sensitive cells assayed by in situ confocal microscopy cellular immunofluorescence staining analysis. WGA (wheat germ agglutinin) as the cell membrane marker (Green). Scale bars, 50 μm. (B) Concentration-effect of sotorasib on H2122 sotorasib-resistant (H2122SR) versus H2122 and H358 sotorasib-tolerant (H358ST2) versus H358 cells. H2122SR and H358ST2 are described in Method section. (C) Western blot analysis of GLUT6 protein expression in H2122SR versus H2122 and H358ST2 versus H358 cells. (D and, E) Abrogation of enhanced glucose uptake and glycolysis in resistant cells by knockdown of GLUT6 assayed in fluorescent glucose analog 2-NBDG-based flow cytometry (D) and Cell Energy Phenotype Test (E) analyses. Cells were transfected with 20 nM siGLUT6 or Mock control for 48 h. The assay details are described in the Method section. (F) Selective inhibitory effect of GLUT6 knockdown on resistant cells versus parental cells in cell growth analysis. Cells were transfected with 20 nM siGLUT6 or Mock control and monitored by IncuCyte ZOOM system every 4 hours. Quantitative data are data are representative of at least 2 independent experiments with 3 technical replicates and presented as mean ± SEM. *P* values were determined using two-way ANOVA followed by Dunnett’s multiple comparisons test. *, *P* < 0.05; **, *P* < 0.01; ***, *P* < 0.001; ****, *P* < 0.0001.

**Figure S4.**
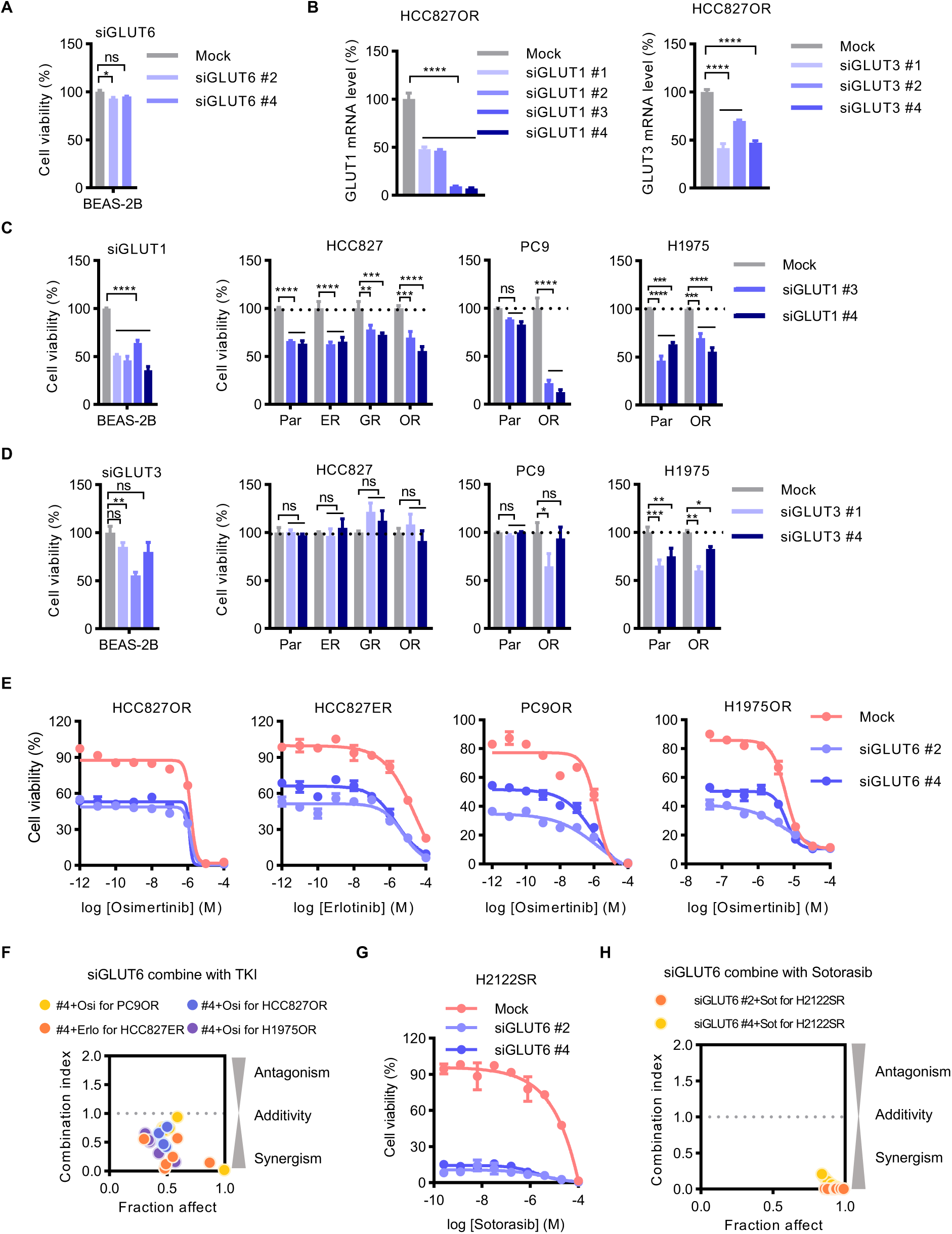
GLUT6 determines resistance to targeted therapy. (A) Negligible effect of GLUT6 knockdown (20 nM siRNA for 72 h) on normal airway epithelial cell line BEAS-2B viability. (B) GLUT1 and GLUT3 mRNA expression after knockdown with siRNA. (C and D) Effect of knockdown of GLUT1 (C, 20 nM siRNA for 72 h) or GLUT3 (D, 20 nM siRNA for 72 h) on the cell viability of the normal airway epithelial cell BEAS-2B and on the various resistant models. (E and F) Synergy effect of GLUT6 knockdown (20 nM for 72 h) on sensitivity to various EGFR inhibitors in resistant cells as shown by drug concentration-effect curves (E) and by Fraction affect - Combination index diagram (F). The Combination index was calculated as indicated in the Methods section. (G and H) Synergy effect of GLUT6 knockdown (20 nM for 72 h) on sensitivity to sotorasib in resistant cells as shown by drug concentration-effect curves (G) and by Fraction affect - Combination index diagram (H). For (B), (C), and (D), data are representative of at least 2 independent experiments with at least 3 technical replicates and presented as means ± SEM. *, P < 0.05; **, P < 0.01; ***, P < 0.001; ****, P < 0.0001; NS, not significant.

**Figure S5.**
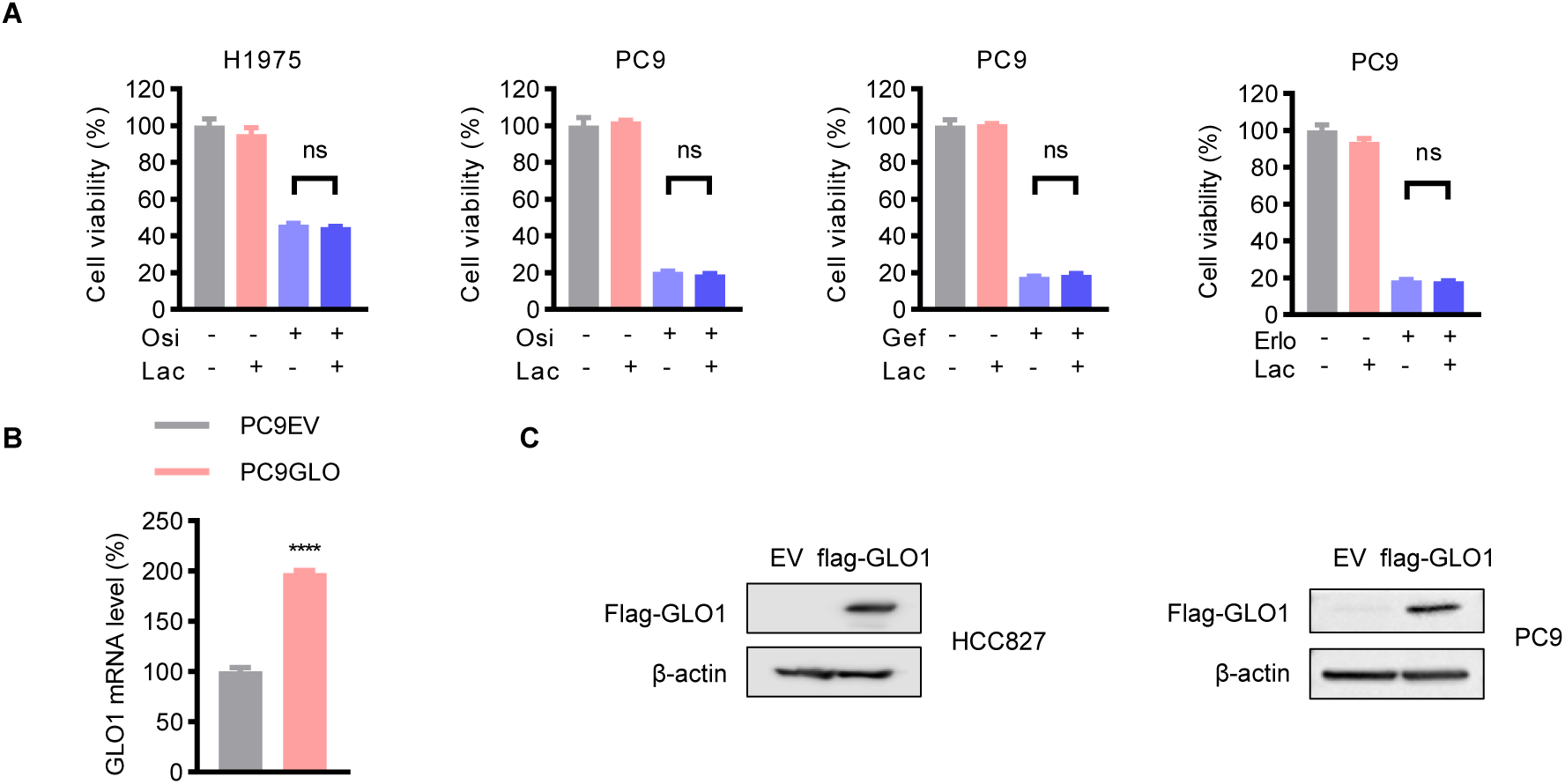
Lactate fails to impart resistance. (A) Negligible effect of lactate (5 mM for 72 h) on EGFRi (500 nM for 72h) sensitivity in cell viability assay. Cells were treated for 72 h. Lac, lactate; Osi, osimertinib; Gef, gefitinib; Erlo, erlotinib. Data are representative of 3 independent experiments with 3 technical replicates and presented as means ± SEM. *P < 0.05, **P < 0.01, ***P < 0.001, ****P < 0.0001; NS, not significant. (B and C) RT-qPCR (B) and western blot (C) analysis of GLO1 expression in GLO1- ectopically-expressed cells. EV, empty vector. β-actin used as loading control.

**Figure S6.**
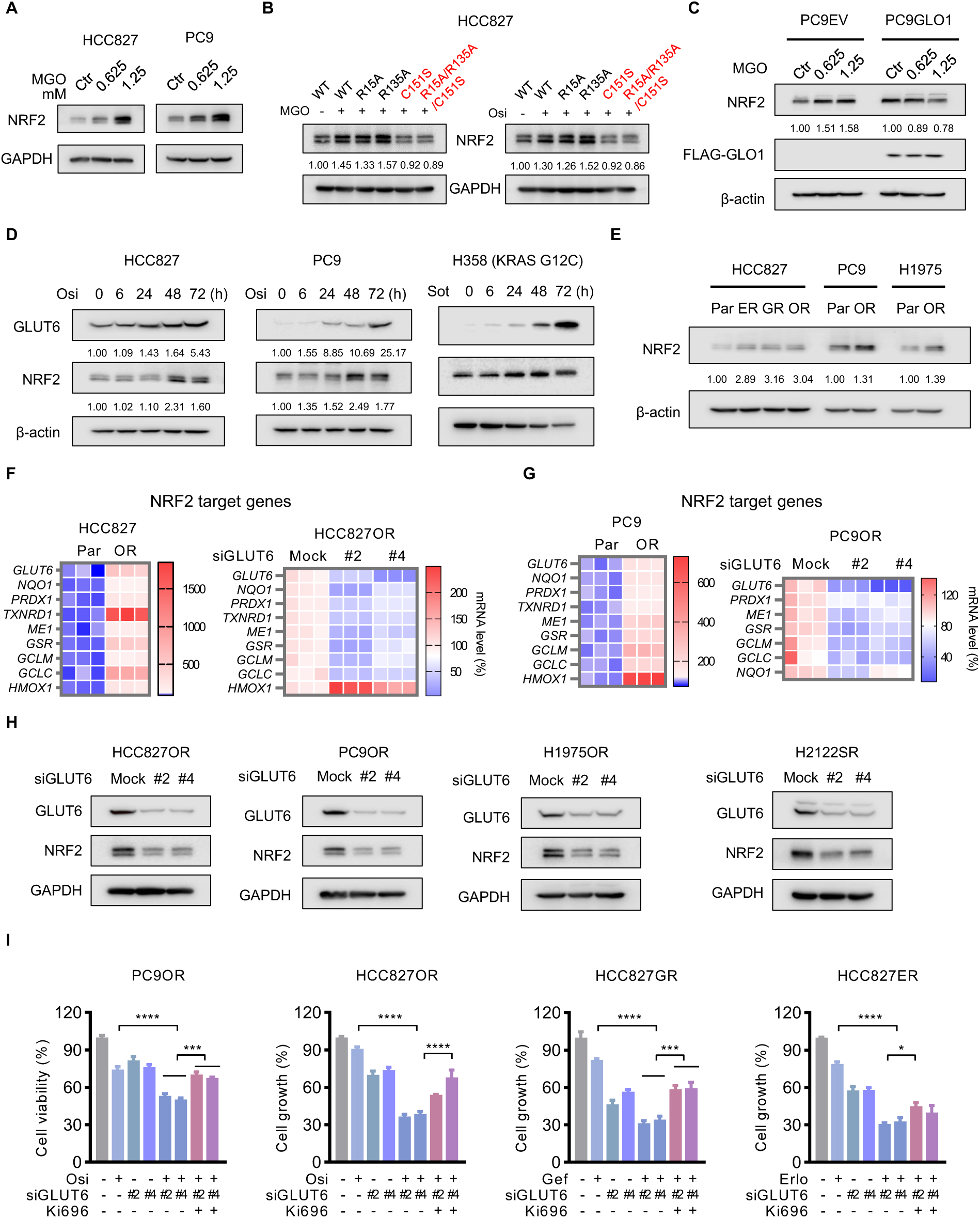
GLUT6-glucose-derived methylglyoxal induces resistance via KEAP1-NRF2 axis. (A) MGO-induced upregulation of NRF2 protein abundance in HCC827 (left panel) and PC9 (right panel) cells assayed by western blot analysis. Cells were exposed to indicated concentrations of MGO for 8 h. (B) MGO (left panel)- and osimertinib (right panel)-induced upregulation of NRF2 protein abundance and its dependence on HMM-formation key amino acid(s) in wild-type (WT) or mutant Flag-KEAP1 expressing HCC827 cells assayed by western blot analysis. Cells were treated by 1.25 mM MGO for 8 h or 500 nM osimertinib for 36 h. (C) Western blot analysis of NRF2 in PC9EV and PC9 Flag-GLO1-ectopically-expressed cells exposed to indicated concentrations of MGO for 8 h. (D) Time-course western blot analysis of the kinetic protein abundance of NRF2 and GLUT6 in HCC827 and PC9 cells treated with 1 µM osimertinib and H358 cells treated with 1 µM sotorasib for indicated periods of time. (E) Western blot analysis of NRF2 protein expression in resistant cells versus their parental counterparts. (F and G) Upregulated expression of NRF2 target gene repertoire in resistant versus parental cells (left panel) and the abrogation effect (right panel) of GLUT6 knockdown (20 nM GLUT6 siRNA for 48 h) assayed by RT-qPCR analysis. (H) Western blot analysis of NRF2 protein expression in EGFRi osimertinib-resistant HCC827OR, PC9OR, and H1975OR cells and KRASi sotorasib-resistant H2122SR cells after GLUT6 knockdown with 20 nM GLUT6 siRNA for 48 h. GAPDH used as loading control. (I) Resensitization of GLUT6 knockdown on osimertinib, gefitinib, and erlotinib in various corresponding resistant cell models and the abrogation effect of NRF2 agonist Ki696. Cells were treated with 5 μM Ki696, 20 nM GLUT6 siRNA, 500 nM EGFR inhibitors, or their combination for 72 h. Data are representative of three independent experiments with three technical replicates and presented as means ± SEM. *, *P* < 0.05; **, *P* < 0.01; ***, *P* < 0.001; ****, *P* < 0.0001.

**Figure S7.**
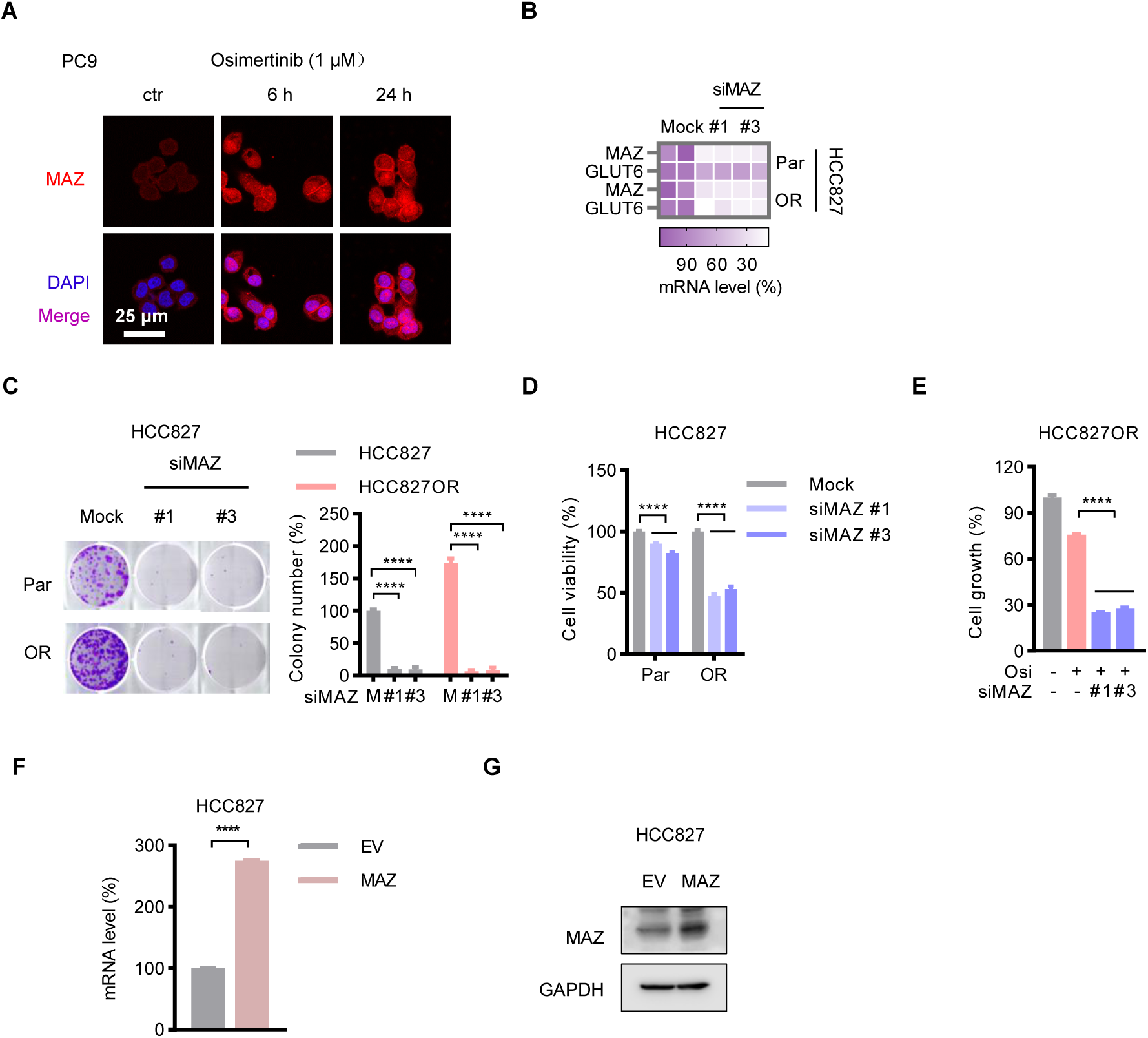
GLUT6 is transcriptionally upregulated by drug stress-activated MAZ. (A) Confocal microscopy immunofluorescence staining analysis shows an increased accumulation of nuclear MAZ (red) induced by osimertinib treatment for indicated periods. Nuclei were counterstained with DAPI (4′,6-diamidino-2-phenylindole). (B) Downregulation of GLUT6 gene expression by knockdown of MAZ with 20 nM siRNA for 24h assayed by RT-qPCR analysis. (C and D) Selective suppression of colony formation (C) and viability (D) with MAZ knockdown in HCC827OR versus parental cells. The cells were transfected with 20 nM MAZ or mock control siRNA for 72 h; in colony formation assay, the cells were resuspended and plated for colony formation for 10 days; in viability analysis, the cells were subject to CCK8 assay at the end of the 72-h treatment. (E) Resensitization of MAZ knockdown (20 nM siRNA for 72 h) to osimertinib in HCC827OR cells. (F and G) RT-qPCR (F) and western blot (G) analyses of MAZ expression in MAZ-ectopically-expressed cells. EV, empty vector. Data are representative of three independent experiments with three technical replicates and presented as means ± SEM. \*\*\**P* < 0.001, \*\*\*\**P* < 0.0001.

**Figure S8.**
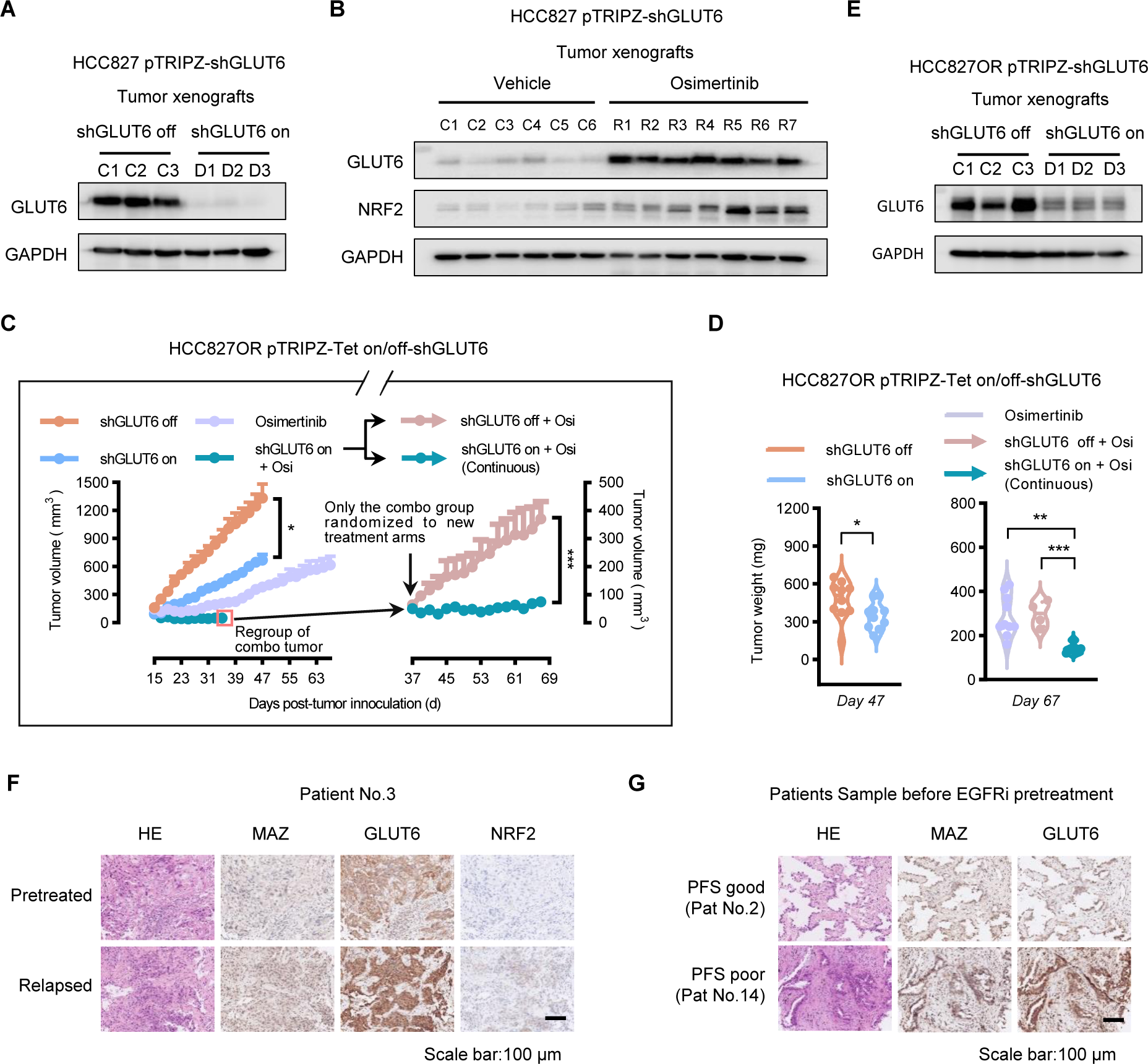
Drug resistance acquisition and maintenance in vivo depend on GLUT6. (A) Western blot analysis of GLUT6 protein expression in HCC827CDX tumors with or without doxycycline-induced knockdown of GLUT6 (shGLUT6 on versus shGLUT6 off). Randomly selected 3 tumor samples per cohort (D1-D3 versus C1-C3, C, control; D, doxycycline treatment.) were subjected to analysis. Tumor information related to Figure 6A, 6B, and 6C. (B) Upregulation of GLUT6 and NRF2 proteins in osimertinib treatment-relapsed tumors assayed by western blot analysis. Tumor information related to Figure 6A, 6B, and 6C; tumor samples in Vehicle (6 tumors) and Osimertinib groups (7 tumors) randomly selected were subjected to analysis. (C and D) Effect of conditional knockdown of GLUT6 with doxycycline-inducible shRNAs (Tet-on-shGLUT6) on overcoming of osimertinib resistance in HCC827ORCDX tumors shown as tumor growth monitoring (C) and tumor weight (D) analyses. Groups: shGLUT6 off (GLUT6 non-knockdown administered with vehicle control, 10 tumors in 5 animals); shGLUT6 on (GLUT6 knockdown induced by Dox, 10 tumors in 5 animals); Osimertinib (10 tumors in 5 animals); and shGLUT6 on + Osi (GLUT6 shRNA + osimertinib, 10 tumors in 5 animals). At day 37 post tumor inoculation, 5 mice in “shGLUT6 on + Osi” group were randomly split and regrouped into shGLUT6 on (continuous) + Osi arm (6 tumors in 3 animals) where the knockdown was continued with Dox administration and shGLUT6 off + Osi arm (4 tumors in 2 animals) where the knockdown was turned off by Dox withdrawal. Vehicle (1% dimethyl sulfoxide, once daily), doxycycline (50 mg/kg, once daily), or osimertinib (2 mg/kg, once daily) was administered through oral gavage. Statistical differences were conducted with repeated measures ANOVA (C) and ANOVA followed by Tukey posttest comparison (D). (E) Western blot analysis of GLUT6 protein expression in HCC827ORCDX tumors with or without doxycycline-induced knockdown of GLUT6 (shGLUT6 on versus shGLUT6 off). Tumor information related to Figure 6A, S8C, and S8D. (F) Upregulation of protein abundance of tumor cell GLUT6 and NRF2, and nuclear MAZ in paired post-treatment resistant and relapsed tumor samples versus pre-treatment tumor samples of the representative 3# patient shown in Figure 6F and 6G assayed in immunohistochemistry (IHC) staining analysis. (G) Upregulation of protein abundance of tumor cell GLUT6 and nuclear MAZ in PFS poor versus PFS good (represented by #14 and #2 patients, respectively, shown in Figure 6H) assayed by IHC staining analysis.

---

**Table S1.**
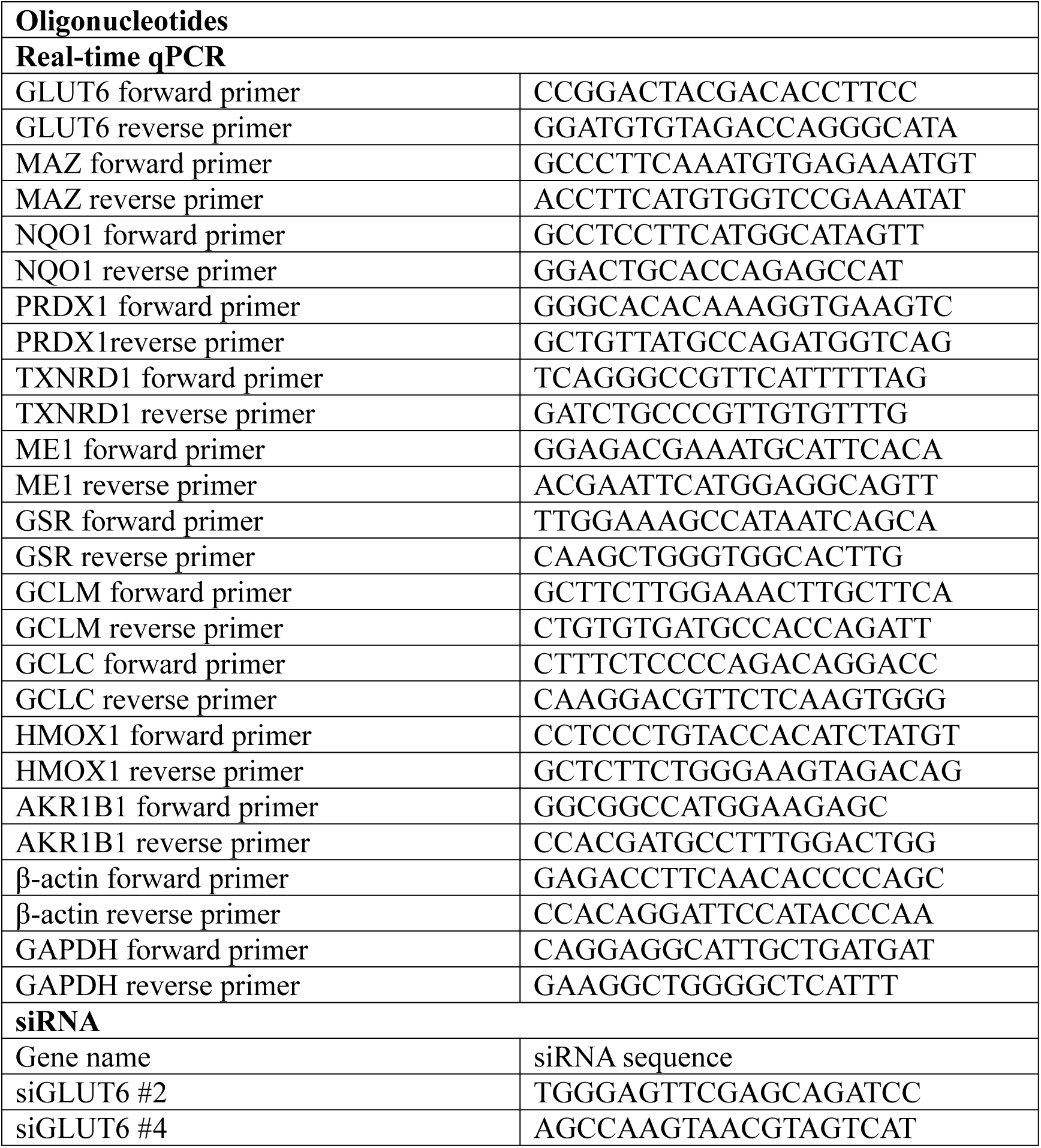
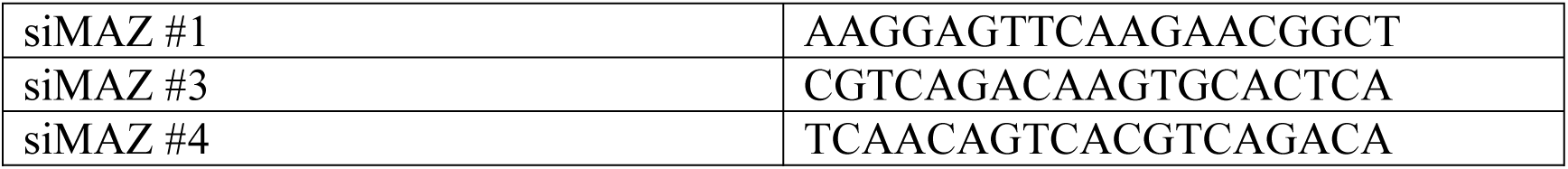
siRNA sequences for knockdown and primers for real-time PCR.

## References

1. Stine, Z.E., Schug, Z.T., Salvino, J.M. & Dang, C.V. Targeting cancer metabolism in the era of precision oncology. Nat Rev Drug Discov 21, 141–162 (2022).

2. Martinez-Reyes, I. & Chandel, N.S. Cancer metabolism: looking forward. Nat Rev Cancer 21, 669–680 (2021).

3. Schmidt, D.R. et al. Metabolomics in cancer research and emerging applications in clinical oncology. CA Cancer J Clin 71, 333–358 (2021).

4. Warburg, O. On the origin of cancer cells. Science 123, 309–314 (1956).

5. Hay, N. Reprogramming glucose metabolism in cancer: can it be exploited for cancer therapy? Nat Rev Cancer 16, 635–649 (2016).

6. Vasan, N., Baselga, J. & Hyman, D.M. A view on drug resistance in cancer. Nature 575, 299–309 (2019).

7. Cooper, A.J., Sequist, L.V. & Lin, J.J. Third-generation EGFR and ALK inhibitors: mechanisms of resistance and management. Nat Rev Clin Oncol 19, 499–514 (2022).

8. Schwenck, J. et al. Advances in PET imaging of cancer. Nat Rev Cancer 23, 474–490 (2023).

9. Vargas, A.J. & Harris, C.C. Biomarker development in the precision medicine era: lung cancer as a case study. Nat Rev Cancer 16, 525–537 (2016).

10. Reck, M. & Rabe, K.F. Precision Diagnosis and Treatment for Advanced Non-Small-Cell Lung Cancer. N Engl J Med 377, 849–861 (2017).

11. Otano, I., Ucero, A.C., Zugazagoitia, J. & Paz-Ares, L. At the crossroads of immunotherapy for oncogene-addicted subsets of NSCLC. Nat Rev Clin Oncol 20, 143–159 (2023).

12. Recondo, G., Facchinetti, F., Olaussen, K.A., Besse, B. & Friboulet, L. Making the first move in EGFR-driven or ALK-driven NSCLC: first-generation or next-generation TKI? Nat Rev Clin Oncol 15, 694–708 (2018).

13. Tan, A.C. & Tan, D.S.W. Targeted Therapies for Lung Cancer Patients With Oncogenic Driver Molecular Alterations. J Clin Oncol 40, 611–625 (2022).

14. Zhang, K.R. et al. Targeting AKR1B1 inhibits glutathione de novo synthesis to overcome acquired resistance to EGFR-targeted therapy in lung cancer. Sci Transl Med 13, eabg6428 (2021).

15. Huang, K. et al. A Novel Allosteric Inhibitor of Phosphoglycerate Mutase 1 Suppresses Growth and Metastasis of Non-Small-Cell Lung Cancer. Cell Metab 30, 1107–1119 e1108 (2019).

16. Luo, M.Y. et al. Metabolic and Nonmetabolic Functions of PSAT1 Coordinate Signaling Cascades to Confer EGFR Inhibitor Resistance and Drive Progression in Lung Adenocarcinoma. Cancer Res 82, 3516–3531 (2022).

17. Lv, Q.M. et al. Cancer cell-autonomous cGAS-STING response confers drug resistance. Cell Chem Biol 30, 591–605 e594 (2023).

18. Deng, D. et al. Molecular basis of ligand recognition and transport by glucose transporters. Nature 526, 391–396 (2015).

19. Echeverria, C., Nualart, F., Ferrada, L., Smith, G.J. & Godoy, A.S. Hexose Transporters in Cancer: From Multifunctionality to Diagnosis and Therapy. Trends Endocrinol Metab 32, 198–211 (2021).

20. Chou, T.C. Theoretical basis, experimental design, and computerized simulation of synergism and antagonism in drug combination studies. Pharmacol Rev 58, 621–681 (2006).

21. Schalkwijk, C.G. & Stehouwer, C.D.A. Methylglyoxal, a Highly Reactive Dicarbonyl Compound, in Diabetes, Its Vascular Complications, and Other Age-Related Diseases. Physiol Rev 100, 407–461 (2020).

22. Bollong, M.J. et al. A metabolite-derived protein modification integrates glycolysis with KEAP1-NRF2 signalling. Nature 562, 600–604 (2018).

23. Schalkwijk, C.G., Micali, L.R. & Wouters, K. Advanced glycation endproducts in diabetes-related macrovascular complications: focus on methylglyoxal. Trends Endocrinol Metab 34, 49–60 (2023).

24. Ma, C.S. et al. NRF2-GPX4/SOD2 axis imparts resistance to EGFR-tyrosine kinase inhibitors in non-small-cell lung cancer cells. Acta Pharmacol Sin 42, 613–623 (2021).

25. Binkley, M.S. et al. KEAP1/NFE2L2 Mutations Predict Lung Cancer Radiation Resistance That Can Be Targeted by Glutaminase Inhibition. Cancer Discov 10, 1826–1841 (2020).

26. Ge, W. et al. iASPP Is an Antioxidative Factor and Drives Cancer Growth and Drug Resistance by Competing with Nrf2 for Keap1 Binding. Cancer Cell 32, 561–573 e566 (2017).

27. LeBoeuf, S.E. et al. Activation of Oxidative Stress Response in Cancer Generates a Druggable Dependency on Exogenous Non-essential Amino Acids. Cell Metab 31, 339–350 e334 (2020).

28. Sayin, V.I. et al. Activation of the NRF2 antioxidant program generates an imbalance in central carbon metabolism in cancer. Elife 6 (2017).

29. Wu, Q. et al. GLUT1 inhibition blocks growth of RB1-positive triple negative breast cancer. Nat Commun 11, 4205 (2020).

30. Nagarajan, A. et al. Paraoxonase 2 Facilitates Pancreatic Cancer Growth and Metastasis by Stimulating GLUT1-Mediated Glucose Transport. Mol Cell 67, 685–701.e686 (2017).

31. Wang, N. et al. Molecular basis for inhibiting human glucose transporters by exofacial inhibitors. Nat Commun 13, 2632 (2022).

32. Byrne, F.L., Olzomer, E.M., Brink, R. & Hoehn, K.L. Knockout of glucose transporter GLUT6 has minimal effects on whole body metabolic physiology in mice. Am J Physiol Endocrinol Metab 315, E286–E293 (2018).

33. Chadt, A. & Al-Hasani, H. Glucose transporters in adipose tissue, liver, and skeletal muscle in metabolic health and disease. Pflugers Arch 472, 1273–1298 (2020).

34. Moraru, A. et al. Elevated Levels of the Reactive Metabolite Methylglyoxal Recapitulate Progression of Type 2 Diabetes. Cell Metab 27, 926–934 e928 (2018).

35. Ferrer, C.M. et al. O-GlcNAcylation regulates cancer metabolism and survival stress signaling via regulation of the HIF-1 pathway. Mol Cell 54, 820–831 (2014).

36. Wang, X. et al. alpha-Ketoglutarate-Activated NF-kappaB Signaling Promotes Compensatory Glucose Uptake and Brain Tumor Development. Mol Cell 76, 148–162 e147 (2019).

37. Cosset, E. et al. Glut3 Addiction Is a Druggable Vulnerability for a Molecularly Defined Subpopulation of Glioblastoma. Cancer Cell 32, 856–868 e855 (2017).

38. Triner, D. et al. Myc-Associated Zinc Finger Protein Regulates the Proinflammatory Response in Colitis and Colon Cancer via STAT3 Signaling. Mol Cell Biol 38 (2018).

39. Ortabozkoyun, H. et al. CRISPR and biochemical screens identify MAZ as a cofactor in CTCF-mediated insulation at Hox clusters. Nat Genet 54, 202–212 (2022).

## References

40. Viswanathan, V.S. et al. Dependency of a therapy-resistant state of cancer cells on a lipid peroxidase pathway. Nature 547, 453–457 (2017).

41. Rees, M.G. et al. Correlating chemical sensitivity and basal gene expression reveals mechanism of action. Nat Chem Biol 12, 109–116 (2016).

